# CXCR4 signaling strength regulates hematopoietic multipotent progenitor fate through extrinsic and intrinsic mechanisms

**DOI:** 10.1101/2023.05.31.542899

**Authors:** Vincent Rondeau, Maria Kalogeraki, Lilian Roland, Zeina Abou Nader, Vanessa Gourhand, Amélie Bonaud, Julia Lemos, Mélanie Khamyath, Clémentine Moulin, Bérénice Schell, Marc Delord, Ghislain Bidaut, Séverine Lecourt, Christelle Freitas, Adrienne Anginot, Nathalie Mazure, David H. McDermott, Véronique Parietti, Niclas Setterblad, Nicolas Dulphy, Françoise Bachelerie, Michel Aurrand-Lions, Daniel Stockholm, Camille Lobry, Philip M. Murphy, Marion Espéli, Stéphane J.C. Mancini, Karl Balabanian

## Abstract

How cell-extrinsic niche-related and cell-intrinsic cues drive lineage specification of hematopoietic multipotent progenitors (MPPs) in the bone marrow (BM) is partly understood. We show that CXCR4 signaling strength regulates localization and fate of MPPs. In mice phenocopying the BM myeloid skewing of patients with WHIM Syndrome (WS), a rare immunodeficiency caused by gain-of-function *CXCR4* mutations, enhanced mTOR signaling and overactive Oxphos metabolism were associated with myeloid rewiring of lymphoid-primed MPPs (or MPP4). Fate decision of MPP4 was also affected by molecular changes established at the MPP1 level. Mutant MPP4 displayed altered BM localization relative to peri-arteriolar structures, suggesting that extrinsic cues contribute to their myeloid skewing. Chronic treatment with CXCR4 antagonist AMD3100 or mTOR inhibitor Rapamycin rescued lymphoid capacities of mutant MPP4, demonstrating a pivotal role for the CXCR4-mTOR axis in regulating MPP4 fate. Our study thus provides mechanistic insights into how CXCR4 signaling regulates the lymphoid potential of MPPs.

## INTRODUCTION

Blood production is a tightly regulated process that starts with hematopoietic stem and progenitor cells (HSPCs). In adults, HSPCs are unique in their capacity to self-renew and replenish the entire blood system through production of a series of increasingly committed progenitor cells within the bone marrow (BM) microenvironment (1, 2). It is now well-established that HSPCs adapt the production of myeloid and lymphoid cells depending on the needs of the body (3). In mice, CD34^-^ long-term (LT)-HSCs form a rare, quiescent population that displays a metabolism skewed towards anaerobic glycolysis at the expense of mitochondrial oxidative phosphorylation (Oxphos) to preserve its quiescent state and long-term reconstitution capacity (4, 5). LT-HSCs differentiate first into CD34^+^ short-term (ST)-HSCs that are characterized by shorter reconstitution ability (6) and then into hematopoietic multipotent progenitors (MPPs), which constitute the stage at which the major divergence of lymphoid and myeloid lineages occurs (7, 8). The MPP compartment can be divided into at least four subsets with different lineage fates (9–11). MPP1 share behaviors exhibited by ST-HSCs including multiple-lineage reconstitution ability (12). By contrast, megakaryocyte/erythroid (ME)-biased MPP2, granulocyte/macrophage (GM)-biased MPP3 and lymphoid-biased MPP4 subsets are more proliferative, devoid of self-renewal potential and likely rely less on glycolysis and more on Oxphos for their metabolism as suggested by transcriptomic analysis (9, 12). The MPP compartment is also dynamic and functionally plastic. Under regenerative conditions, lymphoid-primed MPP4 contribute with their intrinsic GM poising to myeloid output at steady state and undergo a transient change in their molecular identity. This rewires them away from lymphoid differentiation to participate, together with overproduced MPP2 and MPP3, in the burst of myeloid production (10). During ageing, increased rate of myeloid cell production relies on higher number of myeloid-producing HSPCs. The MPP compartment appears to be pivotal in such age-related process as ageing was associated with transcriptomic changes occurring primarily in MPPs, rather than in HSCs, and with a contraction of MPP4 within the BM (13).

The ability of MPPs to express both lymphoid and myeloid specific gene signatures is believed to confer to these cells a capacity to rapidly rewire their differentiation. MPP specification relies on the activity of specific transcription factors (TF), which coordinate expression of pro-lymphoid and -myeloid genes (10, 14–16). Ito-Nakadai and coll. reported that Bach2 and C/EBPβ exert in MPPs opposite effects on the expression of target genes, myeloid genes being activated by C/EBPβ and repressed by Bach2, the latter simultaneously promoting expression of lymphoid genes (15). Recently, deletion of *Ebf1* in all hematopoietic cells has been reported to exacerbate the myeloid potential of MPP3 and MPP4 by enhancing a myeloid signature of chromatin accessibility, overexpressing the myeloid TF *Cebpα* in MPP3 and increasing the capacity of MPP3 and MPP4 cells to generate CD11b^+^ myeloid cells *in vitro* (16). These examples illustrate how activity of different TF tunes MPP commitment into myeloid or lymphoid lineages.

Metabolic plasticity has recently emerged as a critical driver of HSC fate decision (17–19). The metabolite profiles of HSCs and MPPs are similar but differ from restricted progenitor populations and whole BM cells as shown by metabolomics analyses (20, 21). HSCs and MPPs share low levels of glucose uptake, but MPPs display higher mitochondrial volume and membrane potential (22, 23). Metabolomics analyses revealed an enrichment in TCA-cycle related metabolites and amino acids in pooled MPP3/4, while retinoic acid and its downstream product were enriched in HSCs including MPP1 (24). While signaling by retinoic acid appears to play an essential role in maintaining quiescence and self-renewal capacity of HSCs by instructing epigenetic and transcriptional landscapes, it remains to determine whether and how metabolic pathways shape the cellular identity of MPPs.

HSPCs are thought to reside in peri-vascular niches, which regulate their lifelong maintenance and differentiation (25). Peri-sinusoidal stromal cells promote HSC maintenance and retention in the BM notably by secreting niche factors such as Stem cell factor (SCF) and CXCL12 (26–28). Both the localization and stromal niches of the MPP subsets in BM are still incomplete (29). It is also unclear whether regional BM localization of MPP subpopulations impacts their lineage specification and commitment. Although IL-1, IL-6 and TNF-α have been proposed to reprogram MPP4 (30–32), our understanding of how cell-extrinsic niche-related and cell-intrinsic cues drive the lymphoid versus myeloid fate decision of MPPs is still fragmentary. Recently, Kang and coll. reported molecular and functional heterogeneity among MPP3 by identifying a subset of MPP3 that serves as a reservoir for rapid production of myeloid cells in stress and disease conditions (29). This occurs through intrinsic lineage-priming toward myeloid differentiation and cytokine production in the BM. Signaling by the G protein-coupled receptor CXCR4 on MPPs in response to the chemokine CXCL12 produced by BM perivascular stromal cells constitutes a key pathway through which the niche and MPPs communicate. Deletion of *Cxcr4* in total MPPs reduced their differentiation into common lymphoid progenitors (CLPs) and decreased lymphopoiesis (33) but also imbalanced HSC homeostasis as well as access to niche factors such as membrane SCF (34). Using a lymphopenic mouse model bearing a gain-of-function mutation of *Cxcr4* described in the rare human immuno-hematological disorder WHIM Syndrome (WS) (35), we reported that Cxcr4 signaling termination, *i.e.*, desensitization, is required for quiescence/cycling balance of murine HSCs and their differentiation into multipotent and downstream lymphoid-biased progenitors (36). These studies did not address the heterogeneity of lineage-biased MPPs or the molecular mechanisms that promote MPP fate in a Cxcr4-driven manner. Here, we interrogate the mechanisms driving lymphoid-myeloid fate of MPPs by analyzing MPP1-4 carrying or not the gain of function *Cxcr4* mutation. We unveiled that Cxcr4 signaling strength regulates the mitochondrial activity and fate decision of MPP4, while modulating their localization within the BM and thereby likely their access to external cues.

## RESULTS

### The HSPC compartment is myeloid-skewed in the BM of WS mice and patients

To determine whether impaired Cxcr4 desensitization impacted MPP heterogeneity in the BM, a multi-parametric flow cytometry-based strategy was used on the marrow fraction (obtained by centrifugation of the diaphysis) from adult *Cxcr4^+/+^* (WT) and *Cxcr4^1013^*-bearing, *i.e* heterozygous (+/1013) and homozygous (1013/1013), mice as previously reported (9–12). Mutant mice were generated by a knock-in strategy to bear the second most common mutation reported in WS patients at nucleotide 1013, *i.e.*, *CXCR4^1013^*mutation (35). This mutation introduces a stop codon instead of a serine at position 338 in the C-tail of CXCR4 deleting the terminal 15 amino acids and preventing efficient desensitization of the receptor as well as internalization upon activation. MPPs are found in the Lin^-^/Sca-1^+^/c-Kit^+^ (LSK) fraction and the expression levels of CD34, CD48, CD150 and Flt3/CD135 markers divide the MPP pool into MPP1 or ST-HSCs (CD34^+^Flt3^-^CD150^+^CD48^-^), myeloid-biased MPP2 (CD34^+^Flt3^-^ CD150^+^CD48^+^) and MPP3 (CD34^+^Flt3^-^CD150^-^CD48^+^) subsets and the lymphoid-biased MPP4 (CD34^+^Flt3^+^CD150^-^CD48^-/+^) subpopulation (Fig. 1A). First, we assessed the *ex vivo* expression of the signaling trio formed by Cxcl12 and its two receptors Cxcr4 and Ackr3 within MPP subsets. Only Cxcr4 transcripts were readily detectable in all MPPs (Fig. S1A) and Cxcr4 proteins were found at similar levels between WT and mutant subpopulations as determined by flow-cytometric analyses (Figs. 1B and S1B). The absence of Ackr3 and Cxcl12 expression in MPPs was further confirmed in *Ackr3^GFP^*and *Cxcl12^Dsred^* reporter mice (Figs. S1C-E). As expected, +/1013 and 1013/1013 MPPs displayed impaired Cxcr4 internalization following Cxcl12 stimulation as well as increased Cxcl12-promoted chemotaxis in a *Cxcr4^1013^* allele dose- dependent pattern that was abolished by the specific Cxcr4 antagonist AMD3100 (Figs. 1C and S1F). These dysfunctions were not secondary to modulation of membrane Cxcr4 expression but likely relied on the enhanced signaling properties of the truncated Cxcr4 receptor upon Cxcl12 binding as revealed by Akt PhosphoFlow analyses (Fig. 1D). Thus, the desensitization-resistant C-tail-truncated Cxcr4^1013^ receptor is expressed and hyper-functional in all MPP subsets.

**Figure 1:**
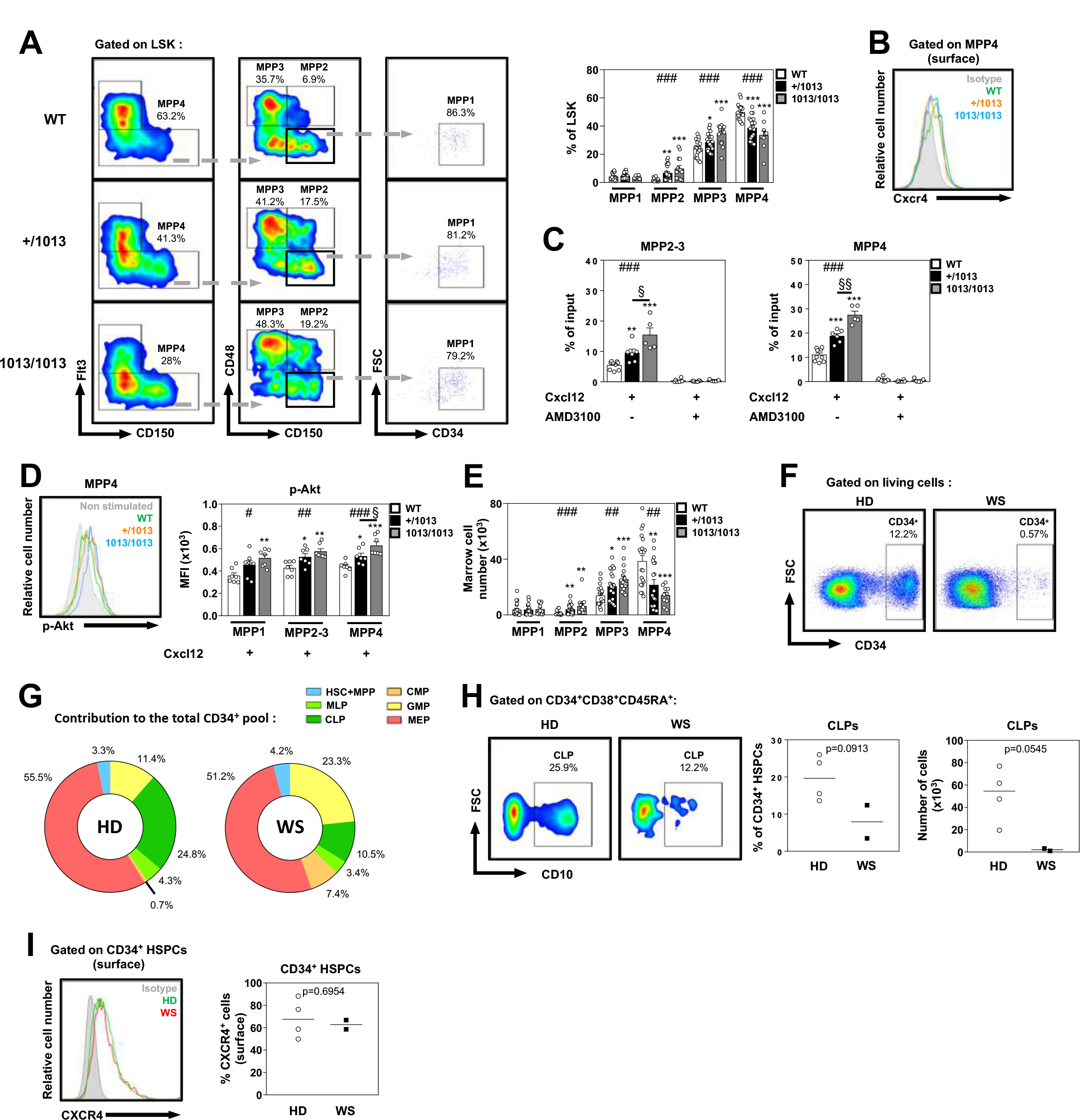
The HSPC compartment is myeloid-skewed in the BM of WS mice and patients. **(A)** Left: Representative flow cytometric analyses comparing frequencies of MPP1 (Lin^-^c- Kit^+^Sca-1^+^ [LSK] CD34^+^Flt3^-^CD150^+^CD48^-^), MPP2 (LSK CD34^+^Flt3^-^CD150^+^CD48^+^), MPP3 (LSK CD34^+^Flt3^-^CD150^-^CD48^+^) and MPP4 (LSK CD34^+^Flt3^+^CD150^-^CD48^+/-^) in the marrow fraction of WT, *Cxcr4^+/1013^* (+/1013) and *Cxcr4^1013/1013^* (1013/1013) mice. Right: Proportions of MPP1, MPP2, MPP3 and MPP4 in the marrow fraction of WT and mutant mice. Data are from at least eight independent experiments with 10-22 mice per group. **(B)** Expression levels of Cxcr4 were determined by flow cytometry. Representative histograms for surface detection of Cxcr4 on gated MPP4 from the marrow fraction of WT and mutant mice. Background fluorescence is shown (isotype, gray histogram). **(C)** Migration of WT and mutant MPP2-3 or MPP4 in response to 10 nM Cxcl12 in presence or absence of 10 µM AMD3100 was assessed by flow cytometry. Data are from three independent experiments with 4-9 mice per group. **(D)** WT and mutant total BM cells were stimulated with 20 nM Cxcl12 at 37°C for 5 min. MPPs were then stained with specific Abs. Left: Representative histograms for intracellular detection of phospho-Akt on gated MPP4 from the marrow fraction of WT and mutant mice. Fluorescence in absence of Cxcl12 stimulation is shown (gray histogram). Right: Mean Fluorescent Intensity (MFI) values for phospho-Akt determined by flow cytometry on gated MPP1, MPP2-3 and MPP4 from the marrow fraction of WT and mutant mice. Data are from three independent experiments with 7 mice per group. **(E)** Absolute numbers of MPP1, MPP2, MPP3 and MPP4 in the marrow fraction of WT and mutant mice. Data are from at least eight independent experiments with 10-22 mice per group. **(F)** Representative dot-plots obtained by flow cytometry comparing the frequencies of CD34^+^ cells in BM aspirates of healthy donors (HD) and WHIM Syndrome (WS) patients. Data are representative of two independent experiments. **(G)** Pie chart representation of the contribution of each HSPC subset to the total CD34^+^ pool in representative HD and WS BM samples. **(H)** Left: Representative dot-plots comparing the frequencies of CLPs (CD34^+^CD38^+^CD45RA^+^CD10^+^). Middle: Proportions of CLPs among CD34^+^ cells. Right: Absolute numbers of CLPs in BM aspirates of HD and WS patients. Horizontal lines correspond to median values. **(I)** Left: Representative histograms for surface detection of CXCR4 on gated CD34^+^ cells from BM aspirates of HD and WS patients. Background fluorescence is shown (isotype, gray histogram). Right: CXCR4-positive fractions obtained within CD34^+^ cells from BM aspirates of HD and WS patients. Horizontal lines correspond to median values. Unless otherwise specified, all displayed results are represented as means ± SEM. Kruskal–Wallis H test–associated p-values (#) are indicated. *, P < 0.05; **, P < 0.005; and ***, P < 0.0005 compared with WT cells; ^§^, P < 0.05; and ^§§^, P < 0.005 compared with +/1013 cells (as determined using the two-tailed Student’s t test). See also Figure S1.

We next compared the BM composition of WT and mutant mice with regards to MPP subsets. We observed a reduction of the proportion and number of MPP4 in a *Cxcr4^1013^* allele dose-dependent pattern (Figs. 1A and E). This was mirrored by an increase of MPP2/3 subsets, thus suggesting that the MPP compartment in *Cxcr4^1013^*-bearing mice is myeloid-biased and defective for lymphoid potential. As previously reported (35, 36), mutant mice had normal numbers of MPP1/ST-HSCs in BM and displayed circulating lympho-neutropenia compared with WT mice (Fig. S1G). Though lineage-biased MPP subsets are not described in the classical human hematopoietic differentiation tree (37, 38), a similar pro-myeloid skewing was observed in BM samples from two unrelated patients with WS who are heterozygous for the autosomal-dominant *CXCR4^R334X^* mutation (Figs. 1F-H and S1H). Frequencies and numbers of CD34^+^ HSPCs tended to decline in the BM of WS patients that encompassed lymphoid-committed progenitors *i.e.*, immature multi-lymphoid progenitors (MLPs; defined as CD45RA^+^CD38^−^CD7^−^CD34^+^CD10^+^) and their progeny CLPs (defined as CD45RA^+^CD38^+^CD34^+^CD10^+^) compared with aged- and sex-matched healthy individuals. Conversely, the frequencies of common myeloid progenitors (CMPs; defined as CD45RA^-^ CD38^+^CD7^−^CD34^+^CD10^-^CD135^+^) and downstream granulocyte-monocyte progenitors (GMPs; defined as CD45RA^+^CD38^+^CD7^−^CD34^+^CD10^-^CD135^+^) tended to increase. Similar surface expression of CXCR4 was observed in healthy and WS HSPCs (Fig. 1I). Thus, these findings unravel a myeloid skewing of the HSPC compartment in the BM of WS mice and patients.

### CXCR4 desensitization is required for efficient generation and maintenance of the lymphoid-biased MPP4 pool

At steady state, MPP subsets are independently produced by ST-HSCs or MPP1 that give rise to higher number of MPP4 than of MPP2 and MPP3 (10). Thus, we hypothesized that the overall myeloid skewing of the mutant MPP compartment could involve an impaired differentiation potential of MPP1. We compared *in vitro* the capacity of MPP1 sorted from the BM of WT or mutant mice to generate biased MPP subsets in feeder-free media (36). By day 4, the number of MPP4 was significantly lower in *Cxcr4^1013^*-bearing cell cultures compared with the WT ones (Fig. 2A). In contrast, mutant MPP1 were as efficient as WT ones at producing MPP2/3 (Fig. 2A and not shown). Accordingly, high-throughput bulk RNA sequencing (RNA-seq) analyses highlighted in mutant MPP1 a gene signature with preserved myeloid potential (Fig. 2B and S2A). Addition of the CXCR4 antagonist AMD3100 rescued the lymphoid potential of mutant MPP1 as it led to an increased number of MPP4, reaching the levels observed in WT cultures (Fig. 2A). Therefore, mutant MPP1 display impaired capacities to differentiate into MPP4 but not into myeloid-biased MPPs and this likely relies on aberrant Cxcr4 signaling. We next interrogated if such imbalance was associated with deregulations in chromatin accessibility by performing Assay for Transposase-Accessible Chromatin sequencing (ATAC-seq) in WT and mutant MPP1. In 1013/1013 MPP1, gene ontology (GO) analyses on regions associated with decreased chromatin accessibility revealed an enrichment for lymphoid differentiation signatures (Fig. 2C and Table S1). To determine whether reduced ATAC-seq peaks harbor TF motifs relevant for lymphoid differentiation, we then performed TF motif enrichment analysis using HOMER. We found within the top 5 highest enriched motifs Fli1, ERG and GABPA TF motifs (Fig. 2D and Table S2). Interestingly, such TF are known to be important regulators of lymphoid differentiation (39–41). These data prompted us to evaluate *in vitro* the ability of WT or mutant MPP1 to generate mature lymphoid cells. When cultured on OP9/IL-7 stromal cells, mutant MPP1 gave rise to approximately 3-times fewer B cells (defined as B220^+^CD11b^-^) compared to WT ones and this was rescued by AMD3100 notably in 1013/1013 cultures (Fig. 2E). Additionally, we assessed using this co-culture system the lymphoid potential of MPP3 and MPP4 generated *in vitro* from MPP1 in feeder-free conditions. As expected, MPP4 sorted from WT MPP1 cultures were more poised to generate B cells than MPP3, indicating that the specific lymphoid skewing of MPP4 was still apparent after feeder-free culture conditions (Fig. 2F). However, mutant MPP4 displayed an impaired capacity to produce B cells, particularly in 1013/1013 cultures, suggesting that Cxcr4-mediated epigenetic changes established at the MPP1 level might be inherited by MPP4 upon differentiation, leading to reduced lymphoid potential at the MPP4 stage. In line with this, we observed by ATAC-seq in 1013/1013 MPP4 an enrichment for lymphoid differentiation signatures in regions associated with decreased chromatin accessibility (Fig. 2G and Table S3). In 1013/1013 MPP4, we found that Fli1 and ERG were among the top 5 highest enriched motifs in reduced ATAC-seq peaks (Fig. 2H and Table S4). Of note, such deregulations were already observed at the MPP1 stage (Fig. 2D and Table S2). Collectively, these findings identify a loss of lymphoid potential at the MPP1 stage in *Cxcr4^1013^*-bearing mice that occurs at the molecular and functional levels and is potentially transmitted to the MPP4 stage upon differentiation.

**Figure 2:**
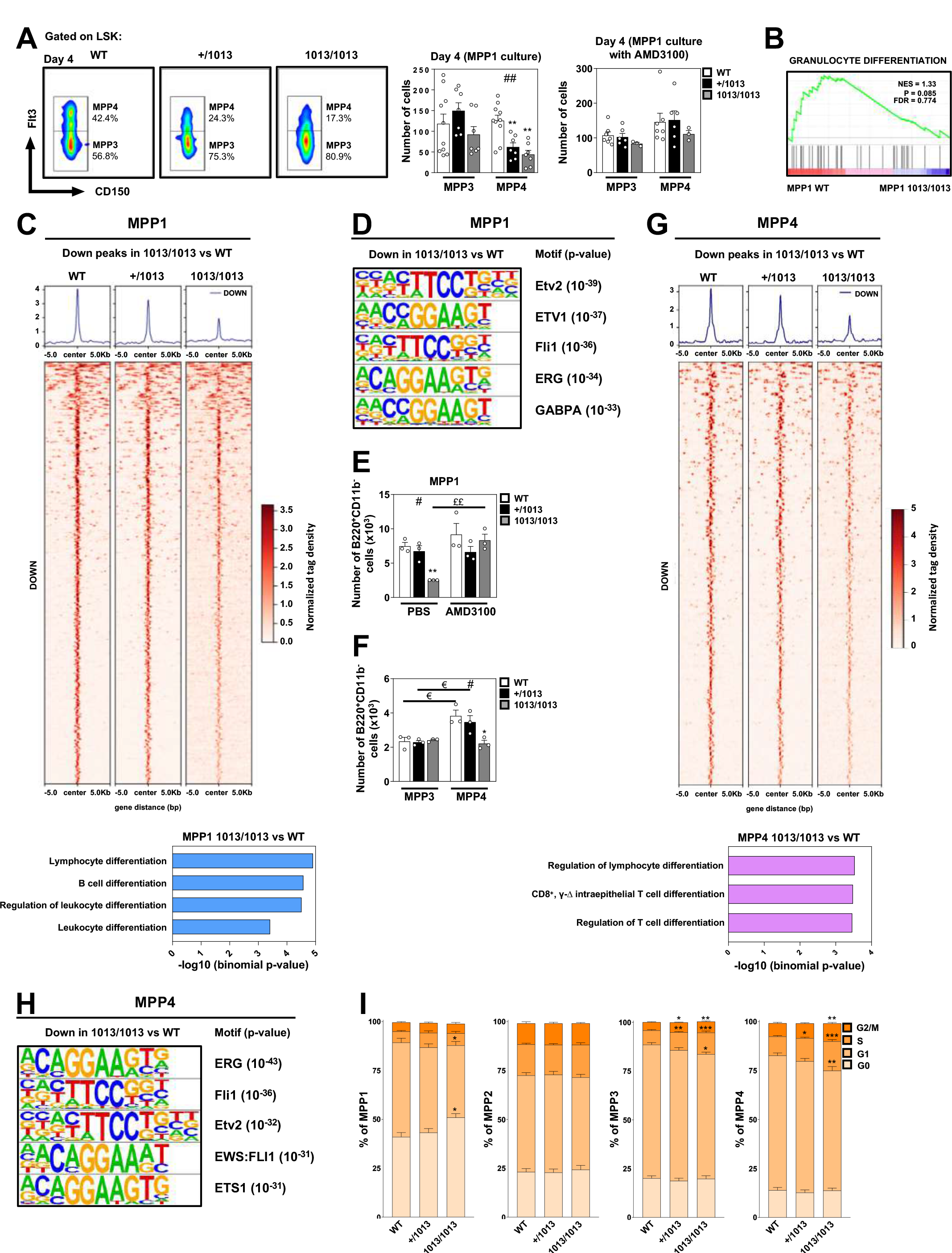
CXCR4 desensitization is required for efficient generation and maintenance of the lymphoid-biased MPP4 pool. **(A)** Equal numbers of ST-HSCs (MPP1) sorted from the marrow fraction of WT or mutant mice were cultured for 4 days in a cytokine-supplemented feeder-free media alone or in the presence of 10 µM AMD3100. Left: Representative flow cytometric analyses comparing the frequencies of MPP3 and MPP4 between WT and mutant MPP1 cultures at day 4 in absence of AMD3100. Right: Absolute numbers of MPP3 and MPP4 in absence or presence of 10 µM AMD3100 determined after 4 days of culture. Data are from four to six independent experiments with 3-10 mice per group. **(B)** RNAseq-based Gene Set Enrichment Analysis (GSEA) for granulocyte differentiation signature in WT vs. 1013/1013 MPP1 is shown. NES = normalized enrichment score, P = nominal p-value, FDR = false discovery rate. **(C)** Upper: Heatmap of down ATAC-seq peaks in WT or mutant MPP1. Lower: Gene ontology analysis with the genes associated with down ATAC-seq peaks in 1013/1013 vs WT MPP1. **(D)** Highest enriched TF motifs in down ATAC-seq peaks in 1013/1013 vs WT MPP1. **(E)** Equal numbers of sorted WT and mutant MPP1 were co-cultured with OP9/IL-7 stromal cells in absence or presence of 10 µM AMD3100. Numbers of B cells were determined after 14 days of culture. Data are from two independent experiments with 3 mice per group. **(F)** MPP3 and MPP4 were sorted from WT or mutant MPP1 after 4 days of culture. Equal numbers of sorted WT and mutant MPP3 or MPP4 were then co-cultured with OP9/IL-7 stromal cells. Number of B cells (B220^+^CD11b^-^) was determined after 7 days of culture. Data are from two independent experiments with 3 mice per group. **(G)** Upper: Heatmap of down ATAC-seq peaks in WT or mutant MPP4. Lower: Gene ontology analysis with the genes associated with down ATAC-seq peaks in 1013/1013 vs WT MPP4. **(H)** Highest enriched TF motifs in down ATAC-seq peaks in 1013/1013 vs WT MPP4. **(I)** Ki-67 and DAPI co-staining was used to analyze by flow cytometry the cell cycle status of MPP1-4 subsets in the marrow fraction of WT and mutant mice. Bar graphs show the percentage of the indicated MPP subsets in G0 (DAPI^low^Ki-67^-^), G1 (DAPI^low^Ki-67^+^), S (DAPI^+^Ki-67^+^) and G2/M (DAPI^high^Ki-67^+^) phases. Data are from five independent experiments with 12-15 mice per group. Unless otherwise specified, all displayed results are represented as means ± SEM. Kruskal–Wallis H test– associated p-values (#) are indicated. *, P < 0.05; **, P < 0.005; and ***, P < 0.0005 compared with WT cells; ^££^, P < 0.005 compared with PBS-treated cells; ^€^, P < 0.05 compared with MPP3 (as determined using the two-tailed Student’s t test). See also Figure S2.

We then investigated by flow cytometry the impact of the *Cxcr4* mutation on both survival and proliferative capacities of MPPs. Abnormal Cxcr4 signaling was associated with decreased fractions of apoptotic cells (Annexin-V^+^DAPI^-^) in *Cxcr4^1013^*-bearing myeloid-biased MPP2/3, while no changes were observed in MPP4 (Fig. S2B). With regards to the cycling status, a significant increase in the frequency of cells in the S/G2-M phases (DAPI^+^Ki-67^+^) was observed among mutant MPP3 and MPP4 subsets (Figs. 2I and S2C). This was associated with reduced frequencies of cells in the G1 phase, thus indicating enhanced G1-S transition in *Cxcr4^1013^*-bearing MPP3 and MPP4. In contrast, the MPP2 pool was unchanged, while mutant

MPP1 displayed increased quiescence as previously reported (36). This was confirmed by *in vivo* BrdU pulse-chase assays. After a 12-days pulse period, we observed a *Cxcr4^1013^* allele dose-dependent increase in BrdU incorporation within mutant MPP3 and MPP4 (Fig. S2D). After 3 weeks of chase, higher proportions of *Cxcr4^1013^*-bearing MPPs had lost label-retaining cell activity compared with WT ones. Therefore, the quiescence/cycling balance and survival profile are differentially disturbed in mutant MPP subpopulations and those dysregulations may contribute to the myeloid skewing by promoting the accumulation of MPP2/3. Paradoxically, increased cycling of mutant MPP4 is associated with contraction of this pool, which evokes aging-induced loss of MPP4 with altered cycling and lymphoid priming (42). Altogether, these results indicate that proper Cxcr4 desensitization is required for efficient generation and maintenance of the lymphoid-biased MPP4 pool.

### CXCR4 desensitization regulates the BM localization of MPP4

To determine whether changes in the MPP compartment of *Cxcr4^1013^*-bearing mice resulted from cell-intrinsic or -extrinsic defects, we performed reciprocal long (16 weeks)- and short (3 weeks)-term BM reconstitution experiments. First, BM cells from WT, +/1013 or 1013/1013 CD45.2^+^ mice were transplanted into lethally irradiated WT CD45.1^+^ recipients (Fig. 3A). Three or sixteen weeks later, there were significantly lower numbers of MPP4 in the BM of *Cxcr4^1013^*-bearing BM-chimeric mice compared with WT chimeras (Fig. 3B). This was mirrored by a slight increase in MPP2/3 subsets. We performed reverse chimeras in which CD45.2^+^ WT and mutant mice were irradiated and reconstituted with WT CD45.1^+^ BM (Fig. 3C). The numbers of MPP4 in the BM were comparable in all groups, whereas MPP2/3 were slightly increased in the BM of mutant mice (Fig. 3D). Together with *in vitro* differentiation findings (Fig. 2A), these results suggest that the contraction in MPP4 in the BM of *Cxcr4^1013^*- bearing mice involves cell-intrinsic effects of impaired Cxcr4 desensitization, while the expansion of myeloid-biased MPPs results from combinatorial effects of aberrant Cxcr4 signaling in both stromal (extrinsic signals) and hematopoietic (intrinsic signals) cells.

**Figure 3:**
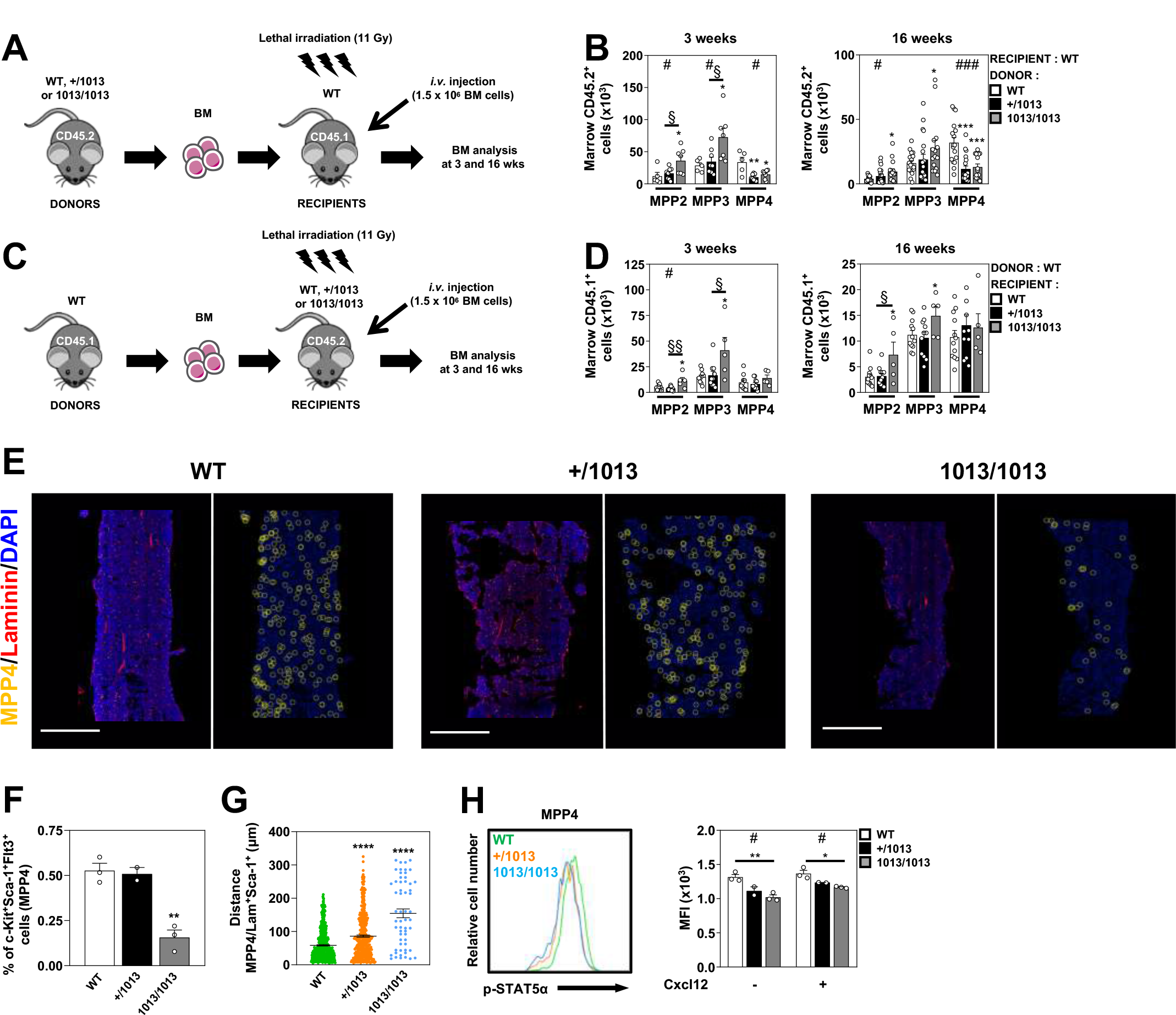
CXCR4 desensitization regulates the BM localization of MPP4. (A and C) Schematic diagrams for the generation of short- and long-term BM chimeras. **(B)** Absolute numbers of donor CD45.2^+^ (WT, +/1013, or 1013/1013) MPP2, MPP3 and MPP4 recovered from the marrow of chimeras in CD45.1^+^ WT recipients 3 weeks (left) or 16 weeks (right) after marrow transplantation. For 3-weeks chimeras, data are from three independent experiments with 5-8 mice per group. For 16-weeks chimeras, data are from four independent experiments with 15-19 mice per group. **(D)** Absolute numbers of donor CD45.1^+^ WT MPP2, MPP3 and MPP4 recovered from the marrow of chimeras in CD45.2^+^ (WT, +/1013, or 1013/1013) recipients 3 weeks (left) or 16 weeks (right) after marrow transplantation. For 3-weeks chimeras, data are from three independent experiments with 5-10 mice per group. For 16-weeks chimeras, data are from four independent experiments with 5-12 mice per group. **(E)** Representative BM sections from WT and mutant mice. MPP4 are indicated in yellow, Laminin in red and DAPI in blue. Scale bars denote 800 µm. Spatial distribution of centroids of MPP4 cells plot as yellow circles within points of DAPI detected cells (blue points) are shown in the embedded pictures. **(F)** Proportions of MPP4 (c-Kit^+^Sca-1^+^Flt3^+^) detected in the BM sections of WT and mutant mice after imaging. Data are from at least 2 independent determinations with 2-3 mice per group. **(G)** Spatial distance analysis between MPP4’s centroid and nearest arteriolar cell’s centroid within representative BM sections from WT and mutant mice. Dots indicate individual MPP4 analyzed for one representative mouse per group with > 58 cells analyzed. Quantification are representative of 3 independent determinations per group. Mann–Whitney U test was used to assess statistical significance (****, P < 0.0001). **(H)** WT and mutant MPP4 were stimulated or not with 20 nM Cxcl12 at 37°C for 2 min. Left: Representative histograms for intracellular detection of phospho-STAT5α on sorted MPP4 from the marrow fraction of WT and mutant mice left untreated. Right: MFI values for phospho-STAT5α were determined by flow cytometry on sorted MPP4 from the marrow fraction of WT and mutant mice. Data are from two independent experiments with 2-3 mice per group. Unless otherwise specified, all displayed results are represented as means ± SEM. Kruskal–Wallis H test–associated p-values (#) are indicated. *, P < 0.05; **, P < 0.005; and ***, P < 0.0005 compared with WT cells; ^§^, P < 0.05; and ^§§^, P < 0.005 compared with +/1013 cells (as determined using the two-tailed Student’s t test). See also Figure S3.

Although the loss of MPP4 in *Cxcr4^1013^*-bearing mice appeared to arise from hematopoietic cell-intrinsic defects, the possibility that enhanced Cxcr4 signaling results in mislocalization of mutant MPP4 within the BM, and ultimately leads to suboptimal access to external cues required for their maintenance and/or fate, remained to be explored. The BM localization of MPP subsets and their supporting stromal niches are still largely incomplete (29, 43). To evaluate the BM positioning of MPP4, we combined *in situ* hybridization assay RNA- scope and immunofluorescence staining (Figs. 3E and S3A). In line with the contraction in mutant MPP4 observed by flow cytometry (Figs. 1A and E), the frequency of c-Kit^+^Sca-1^+^Flt3^+^ cells that are enriched for MPP4 was decreased in the BM of 1013/1013 mice (Figs. 3E and F). No clustering of WT or mutant MPP4 was observed within BM sections (Fig. S3B). We then investigated the positioning of WT and *Cxcr4^1013^*-bearing MPP4 relative to arteriolar cells defined as Laminin^+^Sca-1^+^ (Fig. S3C) (44, 45). Compared to WT mice, mutant mice displayed preserved proportions of arteriolar cells (Fig. S3D). However, mutant MPP4 were localized at greater distances from arteriolar cells compared to WT MPP4 (Fig. 3G). Reduction of phosphorylated STAT5α has been associated with defective positioning of CLPs and pro-B cells near IL-7-positive perivascular stromal cells (33, 46). This prompted us to assess STAT5α status in WT and mutant MPP4 *ex vivo*. Compared to WT MPP4, mutant MPP4 displayed decreased STAT5α phosphorylation in the absence or presence of Cxcl12 (Fig. 3H). Altogether, these findings are suggestive of an altered *in situ* distribution of MPP4 carrying the gain of function *Cxcr4* mutation within BM that could affect access to critical niche factors controlling their maintenance and fate.

### CXCR4 desensitization regulates the molecular identity of MPP4

We next investigated the impact of the gain-of-Cxcr4-function on the molecular identity of biased MPP subsets with a focus on MPP4. RNA-seq analyses of sorted bulk cells yielded robust and reproducible data for all samples with more than 8 x 10^7^ reads *per* sample (Fig. S2A). In total, transcripts corresponding to 20,644 genes were identified. The different MPP subsets clustered distinctly in WT and mutant mice as shown by principal component analysis (PCA) (Fig. S4A). We then applied the gene signatures published by Pietras and collaborators (10) and representing the 50 most highly differentially expressed genes in each MPP subset to our RNA-seq data from WT mice. Gene set enrichment analyses (GSEA) revealed that MPP2-4 signatures were enriched in the corresponding MPP subset, thus validating the robustness of our study (Fig. S4B). Unsupervised analyses of RNA-seq data highlighted 71 genes that were differentially expressed between WT and +/1013 MPP4 compared to 716 genes between WT and 1013/1013 MPP4. Volcano plots of differentially regulated genes indicated that most of these genes were enriched in 1013/1013 MPP4 (Fig. 4A). Only 77 genes were differentially expressed between WT and 1013/1013 MPP3 (Fig. S4C), suggesting that MPP3 and MPP4 do not share similar requirements for the Cxcl12/Cxcr4 signaling axis to regulate their molecular identity. We then performed GO analyses on genes up- or down-regulated in 1013/1013 MPP4 compared to WT ones to determine which biological processes were modulated in mutant MPP4. In line with our proliferation findings (Figs. 2I, S2D and S2C), the MPP4 signature in 1013/1013 mice was enriched for genes related to cell cycle progression and regulation (Fig. 4B and Table S5). Moreover, genes related to Oxphos metabolism appeared to be overrepresented in mutant MPP4. In contrast, expression of key regulators of the lymphoid differentiation decreased in mutant MPP4. Altogether, these results unravel an imbalance in the molecular identity of *Cxcr4^1013^*-bearing MPP4.

**Figure 4:**
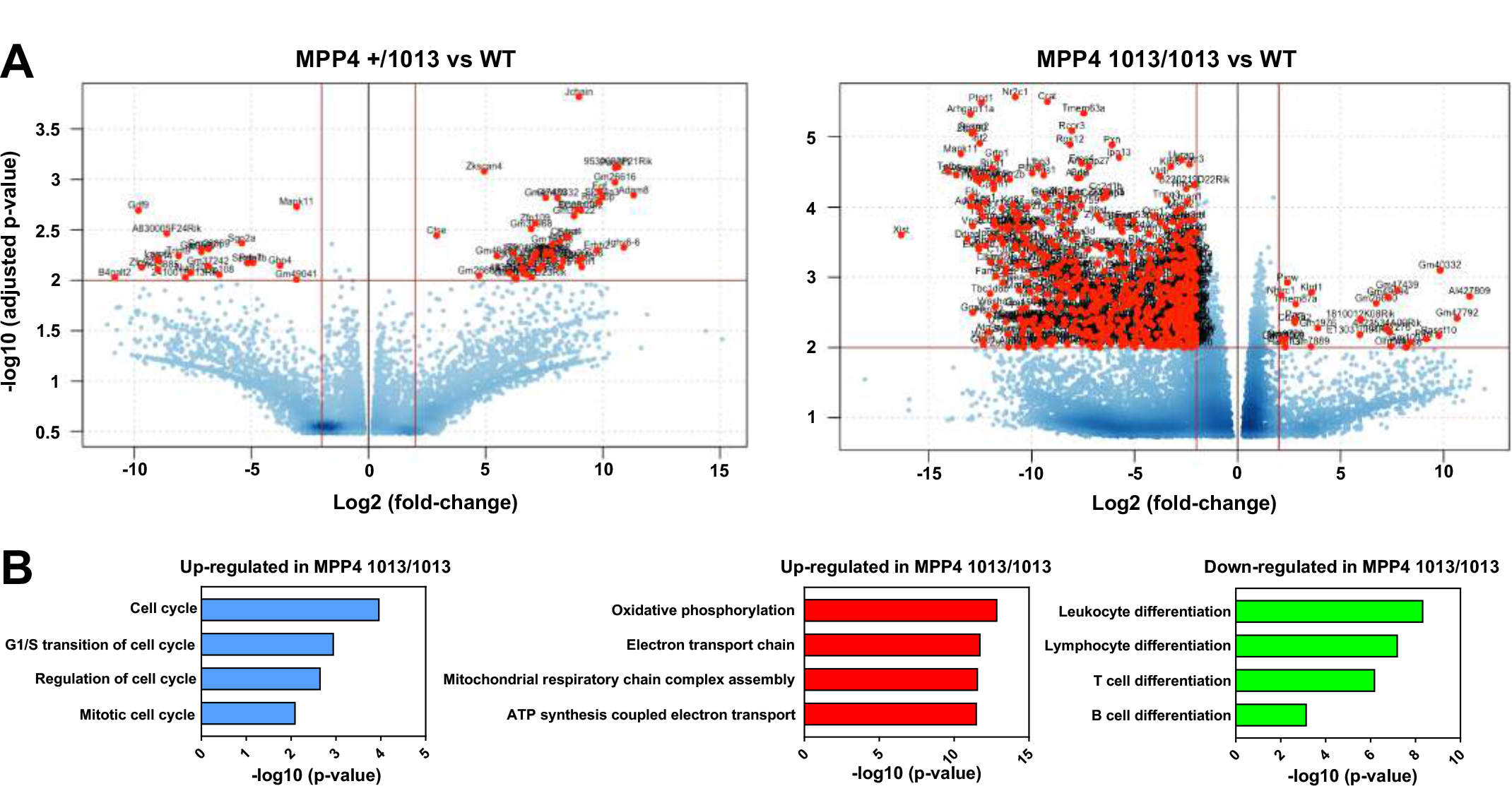
CXCR4 desensitization regulates the molecular identity of MPP4. (A) **Left:** Volcano plot of differentially expressed genes in +/1013 (left) vs. WT (right) MPP4. Right: Volcano plot of differentially expressed genes in 1013/1013 (left) vs. WT (right) MPP4. Genes highly differentially expressed (log2FC(mutant vs WT MPP4) > 2; -log10(adjusted p-value) > 2) are shown in red. Data have been generated by RNA-seq from 3 independent experiments (9 mice for WT and +/1013 MPP4, 6 mice for 1013/1013 MPP4). Three mice of the same genotype have been pooled to generate one replicate. **(B)** Examples of biological processes for which 1013/1013 and WT MPP4 display differential gene expression signatures. Biological processes overrepresented in up- and down-regulated genes in 1013/1013 MPP4 compared to WT MPP4 have been determined by Gene Ontology (GO) analyses. See also Figure S4.

### CXCR4 signaling termination regulates the lymphoid-myeloid gene landscape in MPP4

We determined whether *Cxcr4^1013^*-bearing MPP4 displayed an imbalance in the lymphoid-myeloid gene landscape. Heatmap of RNAseq-based expression data for 48 genes comprised in the GO “Lymphoid differentiation” (GO:0030098) and expressed in WT MPP4 revealed differential expression gene profiles between WT and mutant MPP4 with a significant decrease of the lymphoid signature in 1013/1013 MPP4 (Fig. 5A). This was confirmed by microfluidic-based multiplex gene expression analyses on 0.1 x 10^3^ MPP4. WT and mutant MPP4 clustered differentially along relative expression of master regulators of lymphoid and myeloid differentiation (Fig. 5B and Table S6). There was a significant decreased expression of key lymphoid differentiation genes (*e.g.*, *Ikzf1* and *Dntt*) in mutant MPP4 that followed a *Cxcr4^1013^* allele copy number-dependent pattern (Fig. 5C). This was associated with increased expression of pro-GM genes such as *Mpo* and *Irf8* (47, 48), while the expression of pro-ME genes such as *Gfi1b* (49) was nearly preserved (Fig. 5D). Upregulation of Irf8 in mutant MPP4 was validated at the protein level by flow cytometry (Fig. 5E). Collectively, these findings suggest a global skewing of the MPP compartment that promotes myeloid differentiation likely at the expense of the lymphoid lineage in mutant mice. The fate of the MPP4 subset would be pivotal in such a process.

**Figure 5:**
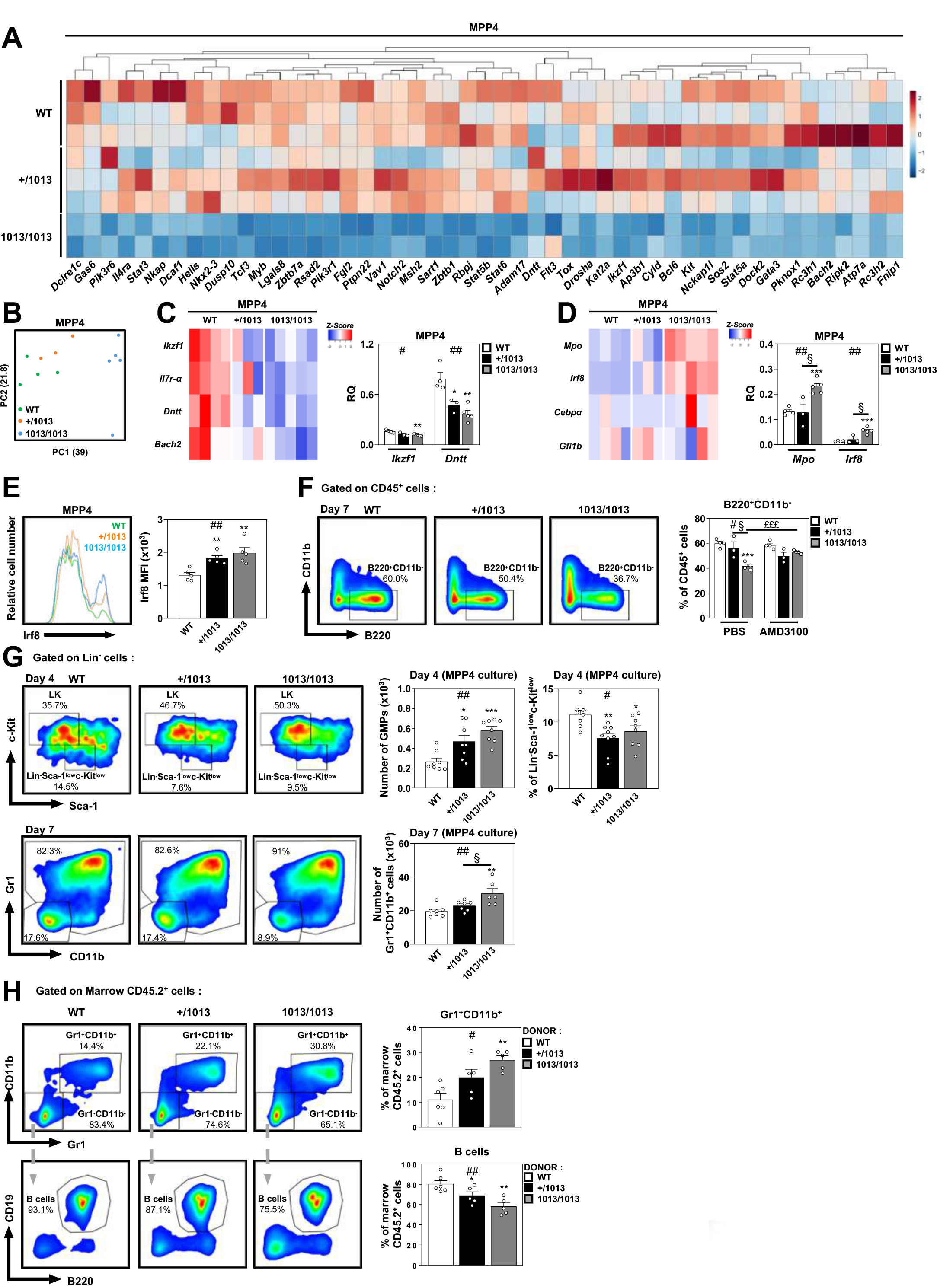
CXCR4 signaling termination regulates the lymphoid-myeloid gene landscape in MPP4. **(A)** RNAseq-based heatmap showing normalized dye intensity for the expression of genes involved in lymphocyte differentiation (GO:0030098) in WT, +/1013 and 1013/1013 MPP4. **(B)** Biomark-based PCA of relative expression of selected genes involved in lympho- myeloid differentiation in MPP4 sorted from WT and mutant mice. **(C)** Left: The heatmap shows the relative quantification (RQ) for lymphoid genes normalized to *Actb* expression levels in each sample (3-5 pools of 0.1 x 10^3^ cells *per* condition). Right: RQ of the most regulated pro-lymphoid genes in WT and mutant MPP4. Data are from two independent experiments with 3-5 mice per group and each individual sample was run in duplicate. **(D)** Left: The heatmap shows the RQ for myeloid genes normalized to *Actb* expression levels in each sample (3-5 pools of 0.1 x 10^3^ cells per condition). Right: RQ of the most regulated pro-GM genes in WT and mutant MPP4. Data are from two independent experiments with 3-5 mice per group and each individual sample was run in duplicate. **(E)** Left: Representative histograms for detection of Irf8 in gated MPP4 from marrow fraction of WT and mutant mice. Right: MFI values for or intracellular Irf8 were determined by flow cytometry in gated MPP4. Data are from two independent experiments with 5 mice per group. **(F)** Equal numbers of sorted WT and mutant MPP4 were co-cultured with OP9/IL-7 stromal cells in absence or presence of 10 µM AMD3100. Left: Representative flow cytometric analyses comparing the frequencies of B cells (B220^+^CD11b^-^) between WT and mutant MPP4 co-cultures at day 7 in absence of AMD3100. Right: Proportions of B cells were determined after 7 days of culture. Data are from two independent experiments with 4 mice per group. **(G)** Equal numbers of sorted WT and mutant MPP4 were cultured for 4 or 7 days in a cytokine-supplemented feeder-free media. Left: Representative flow cytometric analyses comparing the frequencies of Lin^-^Sca-1^-^c-Kit^+^ [LK], Lin^-^Sca-1^low^c-Kit^low^, Gr1^-^CD11b^-^ and Gr1^+^CD11b^+^ cells between WT and mutant MPP4 cultures at day 4 or 7. Middle: Absolute numbers of GMPs were determined after 4 days of culture, while absolute numbers of Gr1^+^CD11b^+^ cells were assessed at day 7. Right: Proportions of Lin^-^Sca-1^low^c-Kit^low^ cells were determined after 4 days of culture. Data are from at least three independent experiments with 6-8 mice per group. **(H)** Proportions of donor CD45.2^+^ (WT, +/1013, or 1013/1013) myeloid cells (Gr1^+^CD11b^+^) or B cells (CD19^+^B220^+^) recovered from the marrow of CD45.1^+^ recipients 2 weeks after the transplantation. Data are from two independent experiments with 5-6 mice per group. All displayed results are represented as means ± SEM. Kruskal–Wallis H test–associated p-values (#) are indicated. *, P < 0.05; **, P < 0.005; and ***, P < 0.0005 compared with WT cells; ^§^, P < 0.05 compared with +/1013 cells; ^£££^, P < 0.0005 compared with PBS-treated cells (as determined using the two-tailed Student’s t test). See also Figure S5.

To test this, we first assessed *in vitro* the B-cell differentiation potential of WT and *Cxcr4^1013^*-bearing MPP4. When cultured on OP9/IL-7 stromal cells, mutant MPP4 differentiation toward B cells was significantly decreased as compared to WT MPP4 and this was rescued by AMD3100 treatment (Fig. 5F). Second, we compared *in vitro* the capacity of 0.5 x 10^3^ MPP4 sorted from the marrow of WT or *Cxcr4^1013^*-bearing mice to generate committed myeloid or lymphoid progenitors and mature cells in feeder-free media. By day 4, both frequencies and numbers of myeloid-committed progenitors including GMPs (defined as LK CD16/32^+^CD34^+^) contained in the Lin^-^Sca-1^-^c-Kit^+^ (LK) fraction were significantly higher in *Cxcr4^1013^*-bearing cell cultures compared with WT ones (Fig. 5G). This likely accounted for the increase in mature myeloid cells expressing both Gr1 and CD11b markers in mutant cultures by day 7. In contrast, the proportions of Lin^-^Sca-1^low^c-Kit^low^ cells that comprise CLPs were reduced in mutant MPP4 cultures compared with the WT ones. Finally, to demonstrate the myeloid bias of *Cxcr4^1013^*-bearing MPP4, we transplanted MPP4 isolated from the BM of WT or mutant CD45.2^+^ mice into sub-lethally irradiated WT CD45.1^+^ recipient mice and analyzed their multi-lineage output 14 days later (Fig. S5A). As expected, WT MPP4 mostly gave rise (∼80%) to lymphoid-derived progeny in the BM of recipient mice (Fig. 5H). In *Cxcr4^1013^*- bearing MPP4-chimeric mice, the lymphoid output was reduced and this was mirrored by increase in donor-derived myeloid cells. A similar increase in myeloid output was observed in the peripheral blood of *Cxcr4^1013^*-bearing MPP4-chimeric mice (Fig. S5B). Taken together, these findings unveil an imbalance in the lymphoid-myeloid gene landscape of *Cxcr4^1013^*- bearing MPP4 that likely affects their capacities to produce lymphoid cells.

### Overactive Oxphos-driven metabolism in *Cxcr4^1013^*-bearing MPP4

Metabolic pathways contribute to HSC self-renewing and fate that rely on differential energy needs (4, 5, 17, 50). When HSCs differentiate, they exit from quiescence and undergo a metabolic switch from anaerobic glycolysis to mitochondrial Oxphos. Transcriptional and metabolomics studies suggest that MPP subsets are more dependent on Oxphos for their metabolism than HSCs, as illustrated by enrichment in TCA cycle-related genes and metabolites compared to HSCs (12, 24). However, whether and how Cxcr4 signaling controls the metabolic profiles of MPP subsets is unknown. GSEA enrichment and heatmap plots of RNA-seq data revealed in mutant MPP4, most particularly in 1013/1013 ones, a significant enrichment in genes encoding for components of Oxphos (Fig. S6A). No alteration of the glycolysis-related gene signature was observed between WT and mutant MPP4 (Fig. S6B).

Flow-cytometric examination of mTOR and downstream targets such as ribosomal protein S6 showed increased phosphorylation in mutant MPP4 in response to Cxcl12 (Fig. 6A), indicating a state of increased metabolic activity including the mTOR pathway. These responses were sensitive to AMD3100 and not observed for the p38 MAPK pathway (Fig. S6C), further suggesting that Cxcl12-promoted Cxcr4 signaling is relayed by the Akt/mTOR pathway in MPP4. In agreement with their overexpressed Oxphos signature, *Cxcr4^1013^*-bearing MPP4 showed increased oxygen consumption rate (OCR), while extracellular acidification rate (ECAR) measurements were reduced compared to WT MPP4 as revealed by Seahorse analyses (Fig. 6B). Oxphos metabolism is associated with the production of mitochondrial metabolites such as reactive oxygen species (ROS). Consistently, mutant MPP4 were poised to produce higher levels of ROS than WT ones as shown by MitoSox, CellRox or DCFDA staining (Fig. S6D). In contrast, mutant MPP4 displayed reduced glucose uptake rate compared to WT MPP4 as shown by 2-NBDG incorporation assay (Fig. S6E). Collectively, these results reveal an overactive Oxphos-driven metabolism in *Cxcr4^1013^*-bearing MPP4.

**Figure 6:**
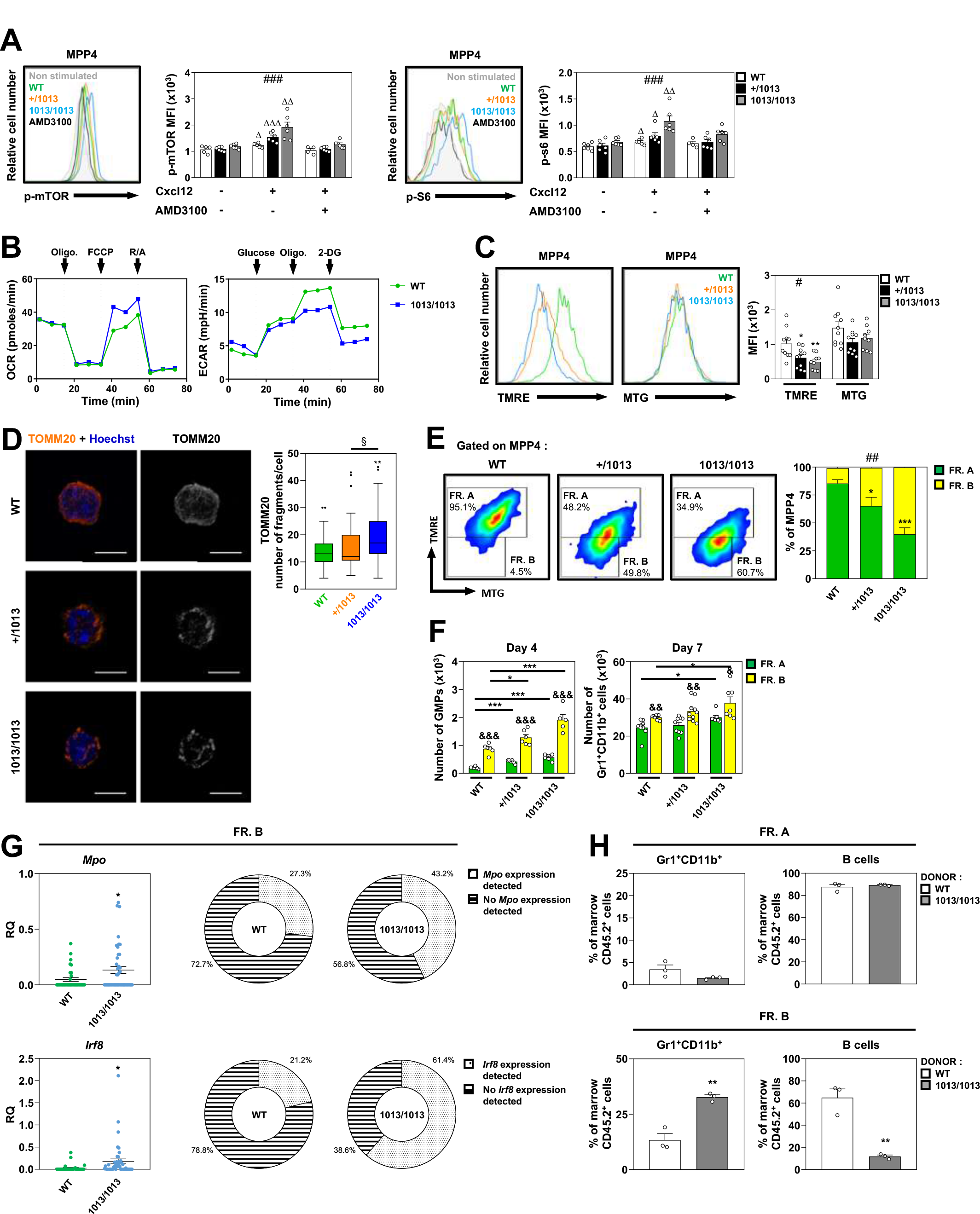
Overactive Oxphos-driven metabolism in *Cxcr4^1013^*-bearing MPP4. (A) **WT and** mutant total BM cells pre-incubated or not with 10 µM of AMD3100 were stimulated or not with 20 nM Cxcl12 at 37°C for 5 min. MPP4 were then stained with specific Abs. Left: Representative histograms for intracellular detection of phospho-mTOR and phospho-S6 on gated MPP4 from the marrow fraction of WT and mutant mice. Fluorescence in absence of Cxcl12 stimulation (gray histogram) or presence of Cxcl12 and AMD3100 (black histogram) are shown. Right: MFI values for phospho-mTOR and phospho-S6 were determined by flow cytometry. Data are from three independent experiments with 4-6 mice per group. **(B)** Oxphos and glycolytic levels measured by oxygen consumption rates (OCR, left) and extracellular acidification rate (ECAR, right) respectively were determined in freshly sorted WT and 1013/1013 MPP4 by Seahorse Mito Stress Test. Curves correspond to the mean of 3 replicates and are representative of one out of three independent experiments. **(C)** Left and middle: Representative histograms for TMRE and MTG staining in gated MPP4 from marrow fraction of WT and mutant mice. Right: MFI values of TMRE and MTG in WT and mutant MPP4. Data are from three independent experiments with 9-10 mice per group. **(D)** Left: Representative staining of TOMM20 in association with Hoechst 33342 in freshly isolated WT or mutant MPP4. Images are representative of three independent determinations. Bars = 5 μm. No staining was observed when the primary Ab was omitted (not shown). Right: Quantitative analyses of number of mitochondrial fragments per cell were determined using FIJI software. Box plots show the median, and cover the interquartile range (IQR) between the first and the third quartiles (Q1 and Q3). Whiskers are drawn at the extreme value that is no more than Q3+1.5xIQR, and no less than Q1-1.5xIQR with outliers shown as black dot. At least 50 cells per condition were analyzed. **(E)** Flow cytometric analyses comparing the frequencies of TMRE^high/+^ fraction A (FR. A) and TMRE^low/-^ fraction B (FR. B) within WT and mutant MPP4. Data are from three independent experiments with 6-8 mice per group. **(F)** Equal numbers of the fraction A or B sorted from WT and mutant MPP4 were cultured for 4 or 7 days in a cytokine-supplemented feeder-free media. Absolute numbers of GMPs were determined after 4 days of culture, while absolute numbers of mature myeloid cells (Gr1^+^CD11b^+^) were assessed at day 7. Data are from three independent experiments with 5-10 mice per group. **(G)** Left: Expression of pro-GM genes in single TMRE^low^ (fraction B) MPP4 from WT and 1013/1013 mice as determined by Biomark. Middle and right: Proportions of WT and 1013/1013 TMRE^low^ MPP4 with detectable or undetectable expression of *Mpo* and *Irf8*. Data are from two independent experiments with >50 cells per condition analyzed. **(H)** Proportions of donor CD45.2^+^ (fraction A or B; WT or 1013/1013) myeloid cells (Gr1^+^CD11b^+^) or B cells (CD19^+^B220^+^) recovered from the marrow of CD45.1^+^ recipients 2 weeks after the transplantation. Data are from two independent experiments with 3 mice per group. All displayed results are represented as means ± SEM. Kruskal–Wallis H test–associated p-values (#) are indicated. *, P < 0.05; **, P < 0.005; and ***, P < 0.0005 compared with WT cells; ^§^, P < 0.05 compared with +/1013 cells; ^Δ^, P < 0.05; ^ΔΔ^, P < 0.005; and ^ΔΔΔ^, P < 0.0005 compared with non-stimulated cells; ^&^, P < 0.05; ^&&^, P < 0.005; and ^&&&^, P < 0.0005 compared with fraction A (as determined using the two-tailed Student’s t test). See also Figure S6.

We next interrogated if these metabolic deregulations were associated with altered mass and activity of mitochondria where Oxphos occurs. WT and mutant MPP4 exhibited similar mitochondrial mass as shown by MitoTrackerGreen (MTG) (19) or mitochondrial transcription factor A (TFAM) (51) staining (Figs. 6C and S6F). Interestingly, decreased levels of mitochondrial membrane potential dye tetramethylrhodamine-ethyl-esther (TMRE) were observed in *Cxcr4^1013^*-bearing MPP4 (Fig. 6C). TMRE is thought to accumulate specifically in active mitochondria and to decrease in damaged mitochondria which are characterized by a loss of membrane potential (19, 52). In agreement, immunofluorescence staining for the translocase of the outer membrane of mitochondria 20 (TOMM20) were punctate in mutant MPP4 as compared to the hyperfused profile in WT ones (Fig. 6D), suggesting the presence of damaged mitochondria in *Cxcr4^1013^*-bearing MPP4. Interestingly, mutant MPP1 also displayed enhanced mTOR signaling and decreased TMRE level (Figs. S6G-I). Therefore, these findings suggest a deteriorated mitochondrial activity in mutant MPP4 that display hyper-proliferative and myeloid-biased status.

Finally, to determine whether the remodeling of mitochondrial machinery in mutant MPP4 was associated with skewing to the myeloid lineage, we sought to segregate the MPP4 subset in two fractions depending on TMRE levels, *i.e.*, positive/high (Fraction A) vs negative/low (Fraction B), enriched in WT and mutant MPP4 respectively (Fig. 6E). Both MPP4 fractions were sorted and characterized for differentiation capacities. *In vitro* feeder-free differentiation assays showed that fraction B from WT MPP4 was more poised to give rise to GMPs and mature myeloid cells than fraction A and this was enhanced from mutant MPP4 (Fig. 6F). These data suggested that increased myeloid output of *Cxcr4^1013^*-bearing MPP4 cultures (Fig. 5G) was likely due to over-representation of the myeloid-biased fraction B at the start of the assay. In agreement, expression levels of key pro-GM genes such as *Mpo* and *Irf8* were increased in fraction B of mutant mice as shown by microfluidic-based qPCR from TMRE^low^ MPP4 at the single cell level (Fig. 6G). Congruent with our *in vitro* differentiation findings (Fig. 6F), WT TMRE^low^ (fraction B) MPP4 were more poised than WT TMRE^high^ (fraction A) MPP4 to give rise to myeloid cells upon transplantation into sub-lethally irradiated WT CD45.1^+^ recipient mice (Fig. 6H). This myeloid output was increased in the BM of 1013/1013 fraction B MPP4-chimeric mice compared to mice transplanted with WT fraction B MPP4. This was mirrored by a decrease in the lymphoid output in the BM of mutant MPP4-chimeric mice. Altogether, these results suggest that accumulation of damaged mitochondria and overactive Oxphos-driven metabolism are associated with myeloid-biased differentiation of MPP4 in absence of Cxcr4 desensitization.

### Modulation of the Cxcr4/mTOR signaling axis normalizes fate properties of *Cxcr4^1013^*- bearing MPP4

The above findings led us to determine whether targeting the Cxcl12/Cxcr4 signaling axis or the mTOR pathway would counteract myeloid skewing of mutant MPP4. First, we evaluated *in vitro* the impact of AMD3100 on the myeloid potential of TMRE^low^ MPP4. Inhibition of Cxcr4 signaling led to a decreased myeloid output by days 4 and 7 from both WT and mutant TMRE^low^ MPP4 cultures (Fig. 7A). This occurred without affecting cell viability and by promoting accumulation of TMRE^low^ MPP4, suggesting a blockage of MPP4 differentiation (Fig. S7A-B). Such a treatment was even sufficient to normalize at day 7 the production of mature myeloid cells from 1013/1013 TMRE^low^ MPP4 cultures, thus supporting a role for Cxcr4 signaling in regulating the myeloid program of MPP4 with altered mitochondrial activity. Then, we assessed the impact of daily intraperitoneal injections for 3 weeks of 5 mg/kg AMD3100 on MPP4 representation in WT and mutant BM (Fig. 7B). Cxcr4 inhibition decreased the number of MPP4 in the BM of WT mice (Fig. 7C). In mutant mice, AMD3100 treatment reversed the quantitative defect in MPP4 with absolute numbers reaching those observed in untreated WT mice. Strikingly, chronic blockade of Cxcr4-dependent signaling corrected the circulating lymphopenia in mutant mice (Fig. 7D). This led us to analyze to what extent AMD3100 treatment impacted the mitochondrial activity of the MPP4 subset. After 3 weeks of treatment, we observed that TMRE levels in mutant MPP4 increased to become similar to WT levels (Fig. 7E). This was due to normalization of the fraction B. Therefore, these data suggest that integrity of the Cxcl12/Cxcr4 signaling pair is required for maintaining the MPP4 subset and its mitochondrial metabolic activity.

**Figure 7:**
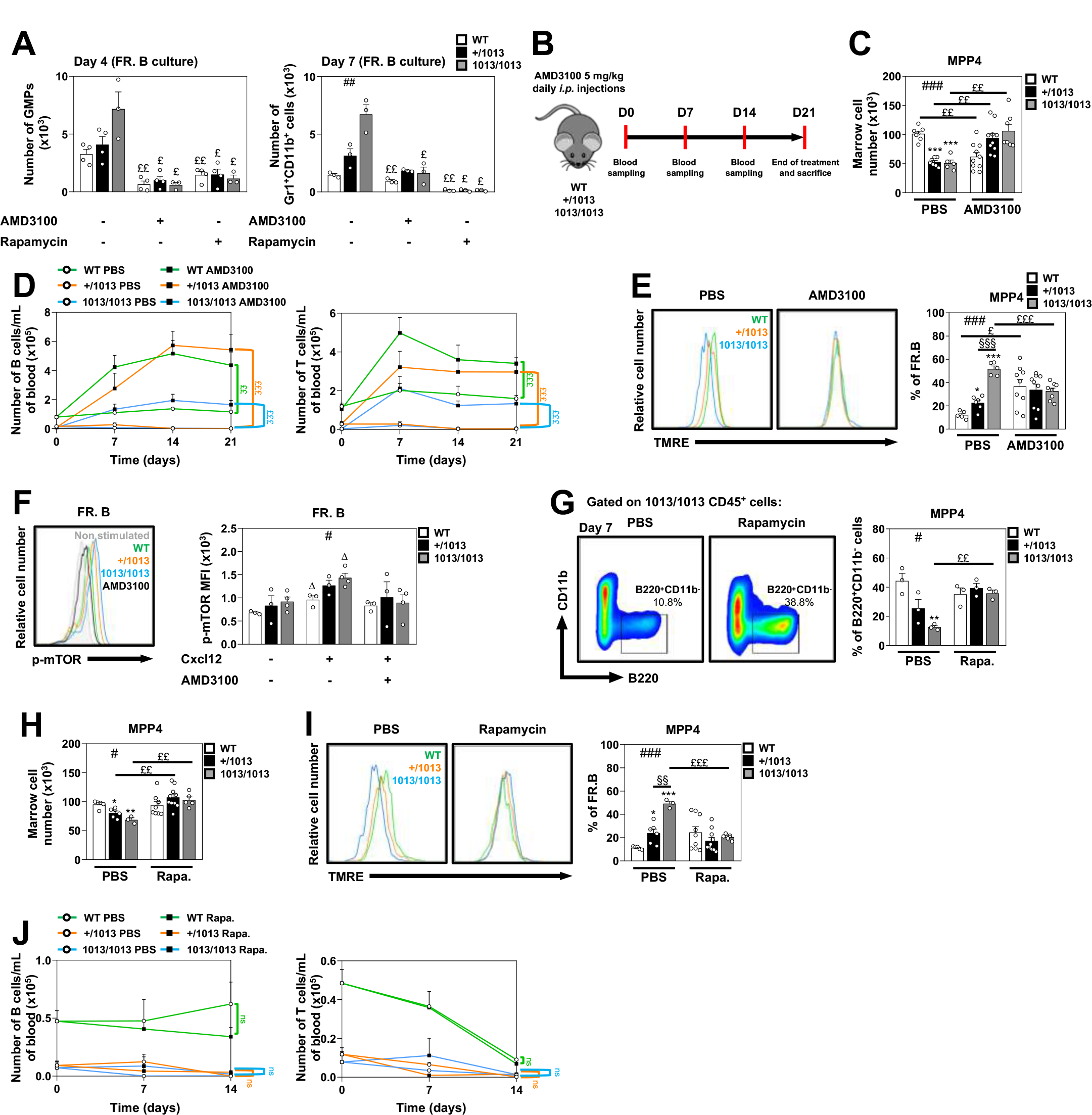
Modulation of the Cxcr4/mTOR signaling axis normalizes fate properties of ***Cxcr4^1013^*-bearing MPP4. (A)** Equal numbers of the fraction B (TMRE^low^) sorted from WT or mutant MPP4 were cultured for 4 or 7 days in a cytokine-supplemented feeder-free media in the presence or absence of 10 µM AMD3100 or 250 nM Rapamycin. Absolute numbers of GMPs were determined after 4 days of culture, while absolute numbers of mature myeloid cells (Gr1^+^CD11b^+^) were assessed at day 7. Data are from two independent experiments with 3-4 mice per group. **(B)** Experimental procedure for *in vivo* AMD3100 treatment. WT, +/1013 and 1013/1013 mice were daily injected intraperitoneally with 5 mg/kg AMD3100 or vehicle (PBS) during 3 weeks. Peripheral blood was drawn and analyzed by flow cytometry at day 0 and then every week. At the end of the treatment, peripheral blood and BM were harvested 2 h after the last injection and analyzed by flow cytometry. **(C)** Absolute numbers of MPP4 were determined by flow cytometry in the marrow fraction of AMD3100-treated WT and mutant mice. Data are from three independent experiments with 5-10 mice per group. **(D)** Numbers of B cells (B220^+^) and T cells (CD3^+^) were determined in the peripheral blood of AMD3100-treated WT and mutant mice every 7 days. Data are from three independent experiments with 5-10 mice per group. **(E)** Left and middle: Representative histograms for TMRE staining in gated MPP4 from AMD3100-treated WT or mutant mice. Right: Proportions of TMRE^low/-^ fraction B among WT or mutant MPP4. Data are from three independent experiments with 5-9 mice per group. **(F)** Fraction B (TMRE^low^) sorted from WT or mutant MPP4 were pre-incubated or not with 10 µM of AMD3100, stimulated or not with 20 nM Cxcl12 at 37°C for 5 min and then stained with specific Abs. Left: Representative histograms for intracellular detection of phospho-mTOR on gated TMRE^low^ MPP4 from the marrow fraction of WT and mutant mice. Fluorescence in absence of Cxcl12 stimulation (gray histogram) or presence of Cxcl12 and AMD3100 (black histogram) are shown. Right: MFI values for phospho-mTOR were determined by flow cytometry in gated TMRE^low^ MPP4. Data are from two independent experiments with 3-4 mice per group. **(G)** Equal numbers of sorted WT and mutant MPP4 were co-cultured with OP9/IL- 7 stromal cells in absence or presence of 250 nM Rapamycin. Left: Representative flow cytometric analyses comparing the frequencies of B cells (B220^+^CD11b^-^) between 1013/1013 MPP4 co-cultures at day 7 in absence or presence of 250 nM Rapamycin. Right: Proportions of B cells were determined after 7 days of culture. Data are from two independent experiments with 3 mice per group. **(H)** Absolute numbers of MPP4 were determined by flow cytometry in the marrow fraction of treated WT and mutant mice 2 h after the last injection of Rapamycin. Data are from two independent experiments with 3-10 mice per group. **(I)** TMRE staining profiles in gated MPP4 from rapamycin-treated WT or mutant mice as described above. Data are from two independent experiments with 3-10 mice per group. **(J)** Numbers of B cells (B220^+^) and T cells (CD3^+^) were determined in the peripheral blood of Rapamycin-treated WT and mutant mice every 7 days. Data are from three independent experiments with 3-10 mice per group. All displayed results are represented as means ± SEM. Kruskal–Wallis H test– associated p-values (#) are indicated. *, P < 0.05; **, P < 0.005 and ***, P < 0.0005 compared with WT cells; ^§§^, P < 0.005 and ^§§§^, P < 0.0005 compared with +/1013 cells; ^£^, P < 0.05; ^££^, P < 0.005; and ^£££^, P < 0.0005 compared with PBS-treated mice; ^Δ^, P < 0.05 compared with non- stimulated cells (as determined using the two-tailed Student’s t test). See also Figure S7.

Finally, we tested the possibility that Cxcr4 signaling regulates metabolic and fate properties of MPP4 through the sensor mTOR. Upon Cxcl12 stimulation *in vitro*, mutant TMRE^low^ MPP4 displayed higher phosphorylated-mTOR content than WT ones and this was abolished by AMD3100 treatment (Fig. 7F). These findings prompted us to evaluate *in vitro* the impact of the mTOR inhibitor Rapamycin on the myeloid potential of TMRE^low^ MPP4. This resulted in accumulation of TMRE^low^ MPP4 and decreased numbers of GMPs and mature myeloid cells from both WT and mutant cultures (Figs. 7A and S7A-B), suggesting an involvement of the mTOR signaling in the fate properties of MPP4. Accordingly, rapamycin treatment restored *in vitro* the capacity of mutant MPP4 to differentiate into B cells when cultured on OP9/IL-7 stromal cells (Fig. 7G). We then determined the consequences of daily intraperitoneal injections for 2 weeks of 4 mg/kg Rapamycin on MPP4 representation in WT and mutant BM. While chronic blockade of mTOR-dependent signaling induced no major changes in WT MPP4, it reversed the quantitative defect in mutant MPP4 and corrected their mitochondrial activity (Figs. 7H and I). Strikingly, rapamycin treatment did not correct the circulating lymphopenia in mutant mice, suggesting that the origin of the lymphopenia is likely multifactorial (Fig. 7J). Collectively, these findings suggest a pivotal role for the Cxcr4-mTOR signaling axis in regulating the fate of MPP4 and their mitochondrial machinery.

## DISCUSSION

This study was designed to determine whether and how CXCR4 proper desensitization controls the lymphoid *versus* myeloid specification of MPPs. We showed that Cxcr4 signaling coordinates the output of the lymphoid and myeloid lineages by regulating the composition and molecular identity of the MPP compartment. The MPP4 subset was pivotal in such a Cxcr4- controlled process. Indeed, Cxcr4 desensitization was required for their efficient generation from MPP1 and then for their maintenance. Cxcr4 signaling seemed also necessary for MPP4 positioning near perivascular niches, and thereby likely for allowing access to external cues that shape their cell cycle status, their metabolism and ultimately their fate. We also provided unanticipated evidence that Cxcr4 signaling regulates the fate decision of MPP4 *via* mTOR.

Impairment of Cxcr4 desensitization resulted in myeloid skewing of the MPP compartment that is characterized by a quantitative defect in lymphoid-primed MPP4 mirrored by an increase in myeloid-biased MPP2 and MPP3. Moreover, analyses of BM samples from two unrelated WS patients carrying the most common WS mutation, *i.e.*, the *CXCR4^R334X^* mutation, unveiled myeloid skewing of the HSPC compartment illustrated by reduced and increased frequencies of lymphoid- and myeloid-committed progenitors, respectively. These findings point towards disturbed HSPC generation and/or differentiation in BM as potential mechanisms contributing to WS-associated lymphopenia (36, 53, 54). This would explain, at least in part, why the lymphopenia can be normalized by chronic AMD3100 treatment in WS mice (this work) and patients (55). However, *in vivo* treatment with Rapamycin did not correct the circulating lymphopenia in mutant mice despite normalizing the number of MPP4 in the BM and restoring their lymphoid potential *in vitro*, suggesting that mechanisms underlying the lymphoid rescue in AMD3100-treated mice might be multifactorial and could involve mobilization of lymphoid cells from peripheral tissues and/or modulation of cytokine production by BM stromal cells. Accordingly, Zehentmeier et *al.* recently reported that daily gavage with the CXCR4 antagonist Mavorixafor restored IL-7 expression and rescued B-cell development in the BM of a WS mouse model bearing the *Cxcr4^R334X^* mutation (56). *Cxcr4^1013^*- bearing mice displayed deregulations of the BM landscape including reduced trabecular and cortical bone structures (57). Interestingly, we reported that intraperitoneal injections of AMD3100 increased the bone mass in *Cxcr4^1013^*-bearing mice by reversing the quantitative defect in skeletal cells and rescuing their osteogenic properties. Whether this partial normalization of the bone environment participates in the rescue of lymphoid compartment in AMD3100-treated mice deserves further investigations.

To date, 29 pathogenic *CXCR4* variants have been identified in WS patients including nonsense, missense and frameshift mutations (58, 59). All of them are positioned in the intracellular region of the C-terminal domain of CXCR4. In all investigated variants, critical phospho-acceptor serine/threonine sites needed for G protein-coupled receptor kinase-mediated phosphorylation are removed and this accounts for the failure of CXCR4 to be desensitized and internalized. These deregulations contribute, together with abnormal β-arrestin–dependent signaling, to enhanced CXCL12-promoted chemotaxis in all investigated variants at the exception of the newly described *CXCR4^Leu371fsX3^* mutation (59, 60). A recent genotype-phenotype correlation study including 14 WS-associated *CXCR4* mutations reported a strong correlation between the degree of internalization defect with the severity of the leukopenia and the recurrence of infections in WS patients (61). Moreover, the CXCR4 antagonist Mavorixafor prevented CXCL12-induced hyperactivation of ERK and AKT *in vitro* in all *CXCR4* variants. Based on these observations, we can speculate that the phenotypes described in our study could be shared, at least in part, with other WS-associated *CXCR4* mutations. In accordance, a new mouse model bearing the *Cxcr4^R334X^* mutation (variant harbored by the two WS patients studied in our work) has been reported recently (56). Although this study did not address the impact of the mutation on early hematopoietic development, the authors showed, in agreement with our results, that this *Cxcr4* gain-of-function mutation was associated with altered B lymphopoiesis in the BM and reduced numbers of naive B cells in blood and secondary lymphoid organs.

Our transcriptomic analyses unveiled a role of Cxcr4 signaling in regulating the molecular identity of MPP4 and particularly their lymphoid-myeloid gene profile. Absence of Cxcr4 desensitization led to reduced accessibility of gene promoters associated with lymphocyte differentiation and decreased expression of key lymphoid differentiation genes in MPP4, while expression of pro-GM genes such as *Mpo* and *Irf8* were increased. Such imbalance in the lymphoid-myeloid gene landscape is likely at the origin of the myeloid skewing of mutant MPP4 we observed at the functional level *in vitro* and *in vivo* after transplantation. Interestingly, the lymphoid potential seemed to be lost as early as the MPP1 stage as shown by impaired capacity of *Cxcr4^1013^*-bearing MPP1 to generate MPP4 and mature lymphoid cells *in vitro*. Accordingly, our ATAC-seq and TF motif enrichment analyses unveiled decreased accessibility of gene promoters and TF motifs associated with lymphocyte differentiation in mutant MPP1. Interestingly, addition of AMD3100 in feeder-free culture conditions rescued the potential of mutant MPP1 to generate MPP4 in absence of Cxcl12. As we did not detect any expression of Cxcl12 by MPP subsets, thus ruling out the possibility of an autocrine signaling of this ligand, these results suggest that mutant MPP1 cultured *in vitro* are still imprinted by the Cxcl12 chemokine they were exposed inside the BM. We also revealed unanticipated evidence that loss of lymphoid potential at the MPP1 stage might be inherited to MPP4. Indeed, we unveiled in mutant MPP4 a decreased accessibility for Fli1 and ERG TF motifs, a deregulation already observed at the MPP1 stage. Moreover, we observed an impaired lymphoid differentiation of MPP4 generated from *Cxcr4^1013^*-bearing MPP1 *in vitro* cultures. Altogether, our findings support a role for the Cxcr4 signaling axis in regulating specification of MPPs at different stages. In such model, the lymphoid potential starts to decrease at the MPP1 stage in mutant mice and this defect is amplified downstream with the myeloid rewiring of the MPP4 subset.

Investigating the BM localization of MPP subsets and their supporting niches remains a challenging issue, mainly due to their scarcity and the absence of specific surface markers. Lai and coll. showed previously that lymphoid priming of MPPs requires the homing of these cells to the inner region of the marrow, a process mediated by G protein-coupled receptor signaling (62). More recently, Cordeiro Gomes and coll. reported that Flt3^+^ MPPs resided in BM niches formed by IL-7^+^ cells comprising peri-sinusoidal stromal cells and endothelial cells (33). Perivascular structures are reported to sustain HSPC maintenance and differentiation through secretion of Cxcl12, SCF and IL-7 (26–28). Here, by combining *in situ* hybridization RNA-scope and immunofluorescence staining, we confirmed the quantitative defect in MPP4 in BM of mutant mice and further showed that proper Cxcr4 signaling is required for MPP4 positioning near peri-arteriolar structures. Reduction of phosphorylated STAT5α has been associated with defective positioning of CLPs and pro-B cells near IL-7^+^ perivascular stromal cells (33, 46). Accordingly, our PhosphoFlow analyses revealed that impairment of Cxcr4 desensitization resulted in decreased STAT5α phosphorylation in MPP4 at baseline, thus supporting the hypothesis that the defective positioning of mutant MPP4 could affect access to critical niche factors controlling their fate such as IL-7. Such mislocalization likely exposed MPP4 to aberrant external cues leading to defects in cell cycle status, mitochondrial activity, maintenance and, ultimately, fate. Recently, Miao *et al.* showed that HSC homeostasis is balanced by the presence of Cxcr4 expressing progenitor cells in the niche, and particularly the MPP4 subset, due to competition for SCF produced by BM stromal cells (34). A major discovery from this study is the displacement of MPP4 and downstream hematopoietic progenitors from Cxcl12-producing niches in the BM through conditional *Cxcr4* deletion. This work points the MPP4 subset as pivotal in regulating HSPC homeostasis within the BM ecosystem by buffering access to extrinsic cues produced by the niches. Recently, Kang and coll. identified a secretory subset of MPP3 that represents a novel bypass mechanism for rapid production of myeloid cells in inflammatory stress and disease conditions (29). This occurs through intrinsic lineage-priming and cytokine production in BM, thereby likely increasing myeloid cell production from other HSPC subsets including MPP4. Whether these changes are observed in the BM of our mouse model and how they would contribute to the lymphoid loss warrant further investigations. Together, these works and the present study illustrate how cell extrinsic and intrinsic mechanisms act together on MPP4 fate and how these processes depend on the integrity of the Cxcl12/Cxcr4 signaling.

Previous transcriptional and metabolomics studies suggested that MPP subsets were more dependent on Oxphos for their metabolism than HSCs. However, whether and how Cxcr4 signaling controls the metabolic profiles of MPP subsets was unknown. Our findings unveiled an overactive Oxphos-driven metabolism in mutant MPP4. This was accompanied by increased ROS production, but counterintuitively, by accumulation of damaged mitochondria with low mitochondrial membrane potential. This metabolically activated cell population was enriched in myeloid-primed cells and was more poised to give rise to myeloid progenies. *In vitro* examination of mTOR in Cxcl12-stimulated mutant MPP4 showed increased phosphorylation at serine 2448, indicating an increase in mTOR activity. This was observed in *Cxcr4^1013^*-bearing MPP4 with low TMRE level as well. Mutant MPP4 also displayed increased phosphorylation of downstream targets such as pS6 at serine 235/236 and Akt at serine 473, supporting the possibility that mutant MPP4 may adopt a state of increased metabolic activity including the mTOR pathway, compatible with high energy production and cycling. Importantly, these responses were sensitive to AMD3100 and Rapamycin, thus suggesting that MPP4 may integrate local cues in BM including Cxcl12 signaling through Cxcr4 and then relayed them by the master regulator mTOR. This would explain why *in vivo* chronic treatment with AMD3100 or Rapamycin normalized fate properties of *Cxcr4^1013^*-bearing MPP4.

Collectively, these findings led us to propose that Cxcr4 acts intrinsically on MPP4 through the mTOR pathway and extrinsically through niche-driven cues to regulate lymphoid cell fate in BM.

## METHODS

### Healthy and WS donors and HSPC analyses

Investigations of human BM samples were performed in compliance with Good Clinical Practices and the Declaration of Helsinki. Cryopreserved BM aspirates from two WS patients (collected under NIH protocol 09-I-0200) were provided through a NIH Material Transfer Agreement in accordance with NIH human subject research policies. None of the patients were under specific treatments aiming at correcting their leukopenia (*i.e.* G-CSF or plerixafor treatments) at the time of the BM collection. BM samples from four healthy donors that were matched for age and sex and used as control subjects were isolated from hip replacement surgery and stored in nitrogen. All individuals have provided written informed consent before sampling and data registering. The study was approved by the Comité de Protection des Personnes Ile-de-France VII (Protocol 17-030, n° ID-RCB: 2017-A01019-44). After sample thawing, BM phenotyping of HSPCs was performed as follows. In brief, 10^6^ cells were labeled for 30 min at 4°C with a combination of the following conjugated Antibodies (Abs): anti– human CD7 (clone M-T701, mouse IgG1), CD10 (clone HI10A, mouse IgG1), CD34 (clone 581, mouse IgG1), CD38 (clone HIT2, mouse IgG1), CD45RA (clone HI100, mouse IgG2b), CD49f (clone GoH3, rat IgG2a), CD135 (clone 4G8, mouse IgG1) and CXCR4 (clone 12G5, mouse IgG2a). Abs were conjugated to brilliant violet (BV) 421, BV605, BV650, PE, PE-cyanin (Cy) 5, peridinin chlorophyll protein PerCP/Cy5.5, alexa fluor (AF) 647 and AF700, and were purchased from BD Biosciences and Biolegend. Corresponding isotype- and species-matched Abs were used as negative controls.

### Mice and genotyping

*Cxcr4^+/1013^* (+/1013) mice were generated by a knock-in strategy and bred as described previously (35). *Cxcr4^1013/1013^* (1013/1013) mice were obtained by crossing heterozygous +/1013 mice. WT mice were used as a control. All mice were littermates and age matched (8– 12 weeks-old). Adult Boy/J (CD45.1) (Charles River) mice were used as transplantation recipients and BM donors. *Cxcl12^Ds-Red^* knock-in mice (*DsRed-Express2* recombined into the *Cxcl12* locus) (27) were obtained from Dr L.G. Ng (Singapore Immunology Network, A*STAR, Singapore) and subsequently crossed with *Cxcr4^1013^*-bearing mice in our animal facility. Transgenic Ackr3-eGFP reporter was provided by Dr. R. Stumm (Institute of Pharmacology and Toxicology, Jena University Hospital, Germany). All the mice were bred in our animal facility under a 12h light/dark cycle, specific pathogen-free conditions and fed ad libitum. All experiments were performed in accordance with the European Union guide for the care and use of laboratory animals and have been reviewed and approved by an appropriate institutional review committee (C2EA-26, Animal Care and Use Committee, Villejuif, France and Comité d’éthique Paris-Nord N°121, Paris, France).

### Sample isolation

BM cells were extracted by centrifugation from intact femurs, tibia and hips to separate marrow and bone fractions. Cell collection was performed in Phosphate-Buffered Saline solution (PBS) 2% Fetal Bovine Serum (FBS) and filtered through a 70-µm nylon strainer to remove debris and fat. All cell numbers were standardized as total count per two legs. The peripheral blood was collected by submandibular puncture. Red blood cell lysis was performed using Ammonium-Chloride-Potassium (ACK) buffer. Analyses were carried out on an LSRII Fortessa flow cytometer (BD Biosciences) and data were analyzed with the FlowJo software (TreeStar).

### Flow cytometry analyses

All staining analyses were performed on an LSR II Fortessa flow cytometer using the following antibodies (Abs): anti–mouse CD3 (clone 145-2C11, hamster IgG1), CD4 (clone RM4-5, rat IgG2a), CD11b (clone M1/70, rat IgG2b), CD16/32 (clone 93, rat IgG2a), CD19 (clone 1D3, rat IgG2a), CD34 (clone RAM34, rat IgG2a), CD41 (clone MWReg30, rat IgG1), CD45R/B220 (clone RA3-6B2, rat IgG2a), CD45.1 (clone A20, mouse IgG2a), CD45.2 (clone 104, mouse IgG2a), CD48 (clone HM48-1, Armenian hamster IgG), CD117 (clone 2B8, rat IgG2b), CD127 (clone A7R34, rat IgG2a), CD135 (clone A2F10, rat IgG2a), CD150 (clone TC15-23F12,2, rat IgG2a), Ter119 (clone TER-119, rat IgG2b), Gr-1 (clone RB6-8C5, rat IgG2b), Sca-1 (clone E13-161,7, rat IgG2a), Irf8 (clone V3GYWCH, mouse IgG1), Ackr3 (clone 8F11-M16, mouse IgG2b) and Cxcr4 (clone 2B11, rat IgG2b). Abs were conjugated to biotin, BV421, BV605, BV650, BV786, FITC, PE, APC, AF700, PE-Cy5, PE-Cy7, eFluor450, AF647, APC-eFluor780, peridinin chlorophyll protein PerCP-Cy5.5, or pacific blue and purchased from BD Biosciences, eBioscience, BioLegend or Sony. The lineage Ab cocktail included anti-CD3, anti-CD45R, anti-CD11b, anti-TER119, anti-CD41, and anti–Gr-1 mAbs (BD Biosciences). Secondary labeling was performed with a Streptavidin–pacific orange or APC-Cy7 from Thermo Fisher Scientific. Data were analyzed with the FlowJo software (TreeStar, Ashland, OR).

### Transplantation experiments and *in vivo* functional assays

For BM transplantation experiments, 1.5 × 10^6^ total marrow cells from young CD45.2^+^ WT, +/1013 or 1013/1013 mice were injected into lethally irradiated young CD45.1^+^ WT recipient mice (two rounds of 5.5 Gy separated by 3 h). For reverse experiments, 1.5 x 10^6^ total marrow cells from CD45.1^+^ WT mice were injected into lethally irradiated CD45.2^+^ WT or mutant recipient mice. Chimerism was analyzed 3 or 16 weeks after reconstitution. For *in vivo* differentiation assay, 10^4^ MPP4 from CD45.2^+^ WT, +/1013 or 1013/1013 mice were injected into sub-lethally irradiated CD45.1^+^ WT recipient mice (one round of 4.5 Gy). BM and blood chimerism were analyzed 2 weeks after reconstitution by flow cytometry. For Cxcr4 blockade experiments, mice were daily injected intraperitoneally with 5 mg/kg AMD3100 (Sigma-Aldrich) or PBS during 3 weeks. Peripheral blood and BM were harvested 2 h after the last injection and analyzed by flow cytometry. For mTOR inhibition experiments, mice were daily injected intraperitoneally with 4 mg/kg Rapamycin (Sigma-Aldrich) or PBS for 2 weeks. Peripheral blood and BM were harvested 2 h after the last injection and analyzed by flow cytometry.

### Cell cycle, survival and metabolic assays

For cell cycle analyses, MPP1-4 from WT and mutant mice were permeabilized and fixed according to the manufacturer’s instructions with the FOXP3 permeabilization kit (Foxp3/Transcription Factor Staining Buffer Set) and then labeled with Ki-67 Ab (clone B56, mouse IgG1; BD Biosciences) and DAPI before flow cytometric analysis. For label-retaining assays (9, 36), mice were injected intraperitoneally with 180 µg BrdU (Sigma-Aldrich) and maintained on water containing 800 µg/ml BrdU and 1% glucose during 12 days. The BrdU pulse was followed by a 3-weeks chase period to detect by flow cytometry label-retaining cells by combining surface staining to define MPPs with intracellular staining using the BrdU-FITC labeling kit following the manufacturer’s instructions (BD Biosciences). Apoptosis was measured using the Annexin V detection kit (BD Biosciences) followed by DAPI staining as described previously (35, 63, 64). For mitochondria analysis, cells were incubated for 20 min at 37°C with tetramethylrhodamine-ethyl-esther (TMRE, 20 nM, Thermo Fisher) and MitoTrackerGreen (MTG, 100 nM, Thermo Fisher) before being analyzed by flow cytometry. Analysis of TFAM was performed as previously described (51) with minor modifications. In brief, 7 x 10^3^ cells were incubated at 4°C for 30 min with surface Abs required to define MPPs before being permeabilized with ice-cold methanol at -20°C for 20 min. After one wash with PBS-2%FBS, cells were incubated with monoclonal rabbit anti-mouse TFAM Ab (Abcam) at room temperature (RT) for 30 min. Cells were then washed, incubated with goat anti-Rabbit IgG Ab at RT for 30 min before being analyzed by flow cytometry.

### Multiplex qPCR

Multiplex qPCR analysis was performed using the microfluidic Biomark system. Bulks of 0.1 x 10^3^ MPP3 or MPP4 or single TMRE^low^ MPP4 were sorted directly into PCR tubes containing 5 µl of reverse transcription/pre-amplification mix as recently described (65). Briefly, the mix contained 2X reaction buffer and SuperScriptIII provided by CellsDirect One-Step qRT–PCR kit (Invitrogen) and 0.2X Taqman assay (Life technologies). Targeted cDNA pre-amplification was performed during 21 (single-cell analyses) or 22 cycles (bulk analyses) and pre-amplified product was diluted 1:5 in TE buffer before processing with Dynamic Array protocol according to the manufacturer’s instructions (Fluidigm). Cells expressing *Actb* and controls genes (*Flt3*, *Cd48*, *Slamf1* and *Irf8*) and not *Pax5* and/or *Cd3* (negatives controls) were considered for further analyses (Table S7). Expression of *Actb* was used for normalization. Heatmaps were generated with http://www.heatmapper.ca (66) using Z scores and principal component analysis (PCA) with R software.

### RNA-seq

For RNA-seq data, pools of 3 x 10^3^ MPP1-4 were sorted directly into RLT buffer (Qiagen) enriched with 1% of β-mercaptoethanol. Total RNA isolation and DNAse treatment were performed using RNeasy Micro Kit (Qiagen). Full length cDNAs were generated from 350 to 1,000 pg of total RNA using Clontech SMART-Seq v4 Ultra Low Input RNA kit for Sequencing (Takara Bio Europe, Saint Germain-en-Laye, France) according to manufacturer’s instructions with 12 cycles of PCR for cDNA amplification by Seq-Amp polymerase. Six hundred pg of pre-amplified cDNAs were then used as input for Tn5 transposon tagmentation by the Nextera XT DNA Library Preparation Kit (96 samples) (Illumina, San Diego, CA) followed by 12 cycles of library amplification. Following purification with Agencourt AMPure XP and SPRIselect beads (Beckman-Coulter, Villepinte, France), the size and concentration of libraries were assessed by capillary electrophoresis. Sequencing reads (50 bp) were generated on the GenomEast platform (Illumina). RNA-seq libraries have been sequenced in Paired-End mode. Briefly, reads were preprocessed in order to remove adapter and low-quality sequences (Phred quality score below 20) and reads shorter than 40 bases were discarded for further analysis. Reads were mapped onto the mm10 assembly of Mus musculus genome using STAR (67). Gene expression quantification was performed from uniquely aligned reads using htseq-count (68) with annotations from Ensembl and “union” mode. Only non-ambiguously assigned reads have been retained for further analyses. Read counts have been normalized across samples with the median-of-ratios method proposed by Anders and Huber (69), to make these counts comparable between samples. Genes expressed in at least 2 biological replicates were considered for further analyses, while 10% of genes with the lowest expression in WT and mutant MPPs summed up were excluded. PCA, volcano plots and GSEA analyses have been generated using R software. GO analyses have been performed using https://biit.cs.ut.ee/gprofiler (70). Data are available in GEO: GSE182992.

### ATAC-seq

ATAC-seq was performed as previously described (71) with some modifications. Briefly, 10^4^ to 3 x 10^4^ cells were spun at 500g for 5 min, washed with cold PBS, lysed in cold lysis buffer (10 mM Tris-HCl, pH 7.5, 10 mM NaCl, 3 mM MgCl2 and 0.1% NP-40, 0.1% Tween-20 and 0.01% Digitonin), incubated 3 min on ice, resuspended in wash buffer (10 mM Tris-HCl, pH 7.5, 10 mM NaCl, 3 mM MgCl2, 0.1% Tween-20) and spun at 500g for 10 min at 4°C. The pelleted nuclei were resuspended in 50 µl transposase reaction mix (1X Diagenode Tagmentation buffer (Diagenode C01019043), 0,1% Tween-20, 0.01% Digitonin, 1.7ug Tn5 (Diagenode C01070010) previously loaded with nextera adapters loaded) for 30 min at 37°C on thermomixer shaking at 1000 rpm. Transposed DNA was purified using a DNA Clean & Concentrator-5 kit (Zymo Research) in 10 µl nuclease-free H2O and amplified with NEBNext® High-Fidelity 2X PCR Master Mix and 1 ul of primer pair from Diagenode (24 SI for tagmented libraries, C01011032), using the following PCR conditions: 72°C for 5 min; 98°C for 30 s then 12 cycles of 98°C for 10 s, 63°C for 30 s and 72°C for 1 min. Libraries were purified and size selected (1.4X-0.5X-1X successively) with AMPure XP beads (Beckman Coulter) then subjected to high-throughput paired-end sequencing (75bp) using the Illumina NextSeq500 sequencer (Illumina, San Diego, CA). Alignment, quality control, peak calling and reproducibility were assessed using ENCODE ATAC-seq analysis pipeline (https://github.com/ENCODE-DCC/atac-seq-pipeline). Briefly, sequencing reads were aligned to the mouse mm10 version of the genome using Bowtie2 (72), duplicate reads marked with Picard Tools and removed, and accessibility peaks were called using MACS2 (73) using default options. BigWig files normalized for the depth of sequencing of aligned filtered reads were generated using deeptools (74). Differential enrichment analysis was performed using diffReps (75) on replicated data using negative binomial statistical test using following parameters: -- meth nb --mode p --nsd s --frag 250. Regions with an abs(log2FoldChange) ≥ 0.8 and an adjusted p-value ≤ 0.001 were kept for downstream analysis. Differential regions annotation and Gene Ontology analysis was performed using GREAT software (http://great.stanford.edu/public/html/) (76). Motif enrichment analysis was performed using Homer v4.11 (77) (findMotifsGenome.pl) on ATAC-seq peaks significantly downregulated (log2FC >= 0.8, pvalue <= 10-5) in 1013/1013 MPP1 or MPP4 compared to WT using the following parameters: mm10 -size 600.

### *In vitro* functional assays

Cxcr4 internalization assay was performed as previously described (35, 36) with some minor modifications. In brief, 1 x 10^6^ BM cells were incubated at 37°C or 4°C for 45 min with or without Cxcl12 (10 nM, R&D Systems). The reaction was stopped by adding ice-cold RPMI and brief centrifugation at 4°C. After one wash in acidic glycine buffer, pH 4.3, levels of Cxcr4 membrane expression were determined by flow cytometry using the additional markers CD3, CD45R, Gr1, CD11b, Ter119, CD41, CD117, Sca-1, Flt3, CD48 and CD150 antigens.

Background fluorescence was evaluated using the corresponding PE-conjugated immunoglobulin-isotype control Ab. Cxcr4 expression in stimulated cells was calculated as follows: (Cxcr4 geometric MFI of treated cells/Cxcr4 geometric MFI of unstimulated cells) × 100; 100% correspond to receptor expression at the surface of cells incubated in medium alone. For chemotaxis assays, preliminary enrichment in Lin^-^ cells was performed using a HSPC enrichment set (BD Biosciences). 1 x 10^6^ of cells were then diluted in 150 µl of DMEM supplemented with 0.5% BSA and 10 mM HEPES and incubated for 45 min at 37°C before being allowed to migrate through a 6.5 mm diameter, 5 µm pore polycarbonate Transwell culture insert (Corning). The same media (450 µl) with or without 10 nM of Cxcl12 was placed in the lower chamber. When specified, the Cxcr4 antagonist AMD3100 (10 µM, Sigma-Aldrich) was added in both upper and lower chambers. Input cells that migrated to the lower chamber after 3 h of incubation at 37°C in humidified air with 5% CO_2_ were collected, stained with CD3, B220, Gr1, Ter119, CD117, Sca-1, CD48, CD150 and Flt3 Abs and counted by flow cytometry. The fraction of cells migrating across the polycarbonate membrane was calculated as follows: [(number of cells migrating to the lower chamber in response to chemokine or medium) / (number of cells added to the upper chamber at the start of the assay)] x 100. For Phosphoflow analyses of mTOR, Akt and S6, 7 x 10^6^ of total BM cells were pre-incubated in DMEM only with or without 10 µM of AMD3100 during 1 h before being stimulated with 20 nM Cxcl12 during 5 min at 37°C. Cells were then stained with CD3, B220, Gr1, Ter119, CD117, Sca-1, CD48, CD150 and Flt3 Abs. Phosphoflow stainings have been performed using the Perfix EXPOSE Kit (Beckman-Coulter) according to the manufacturer’s instructions and Phospho-mTOR (Ser2448), Phospho-Akt (S473) and Phospho-S6 (Ser235/236) Abs. For Phosphoflow analyses of p38 and STAT5α, sorted 10^3^ MPP4 were pre-incubated in DMEM only with or without 10 µM of AMD3100 during 1 h before being stimulated with 20 nM Cxcl12 during 2 min at 37°C. Phosphoflow stainings have been performed using the Perfix EXPOSE Kit as previously and Phospho-p38 (Thr180/Tyr182) and Phospho-STAT5α (Tyr694) Abs. For *in vitro* hematopoietic differentiation experiments, sorted ST-HSCs/MPP1, MPP3 or MPP4 were seeded at 0.5 x 10^3^ cells into a 96-well tissue culture plate in DMEM medium supplemented with 10% FBS, 1% penicillin/streptomycin, and the following murine cytokines (5, 10, or 50 ng/mL): SCF, IL-3, IL-6, IL-7, and Flt3-L (PeproTech) in the presence or absence of 10 µM AMD3100 or 250 nM Rapamycin. Cells were cultured at 37°C in a humidified atmosphere containing 5% CO_2_ and were harvested at the indicated intervals to be analyzed by flow cytometry after staining with CD3, B220, Gr1, CD11b, Ter119, CD117, Sca-1, CD48, CD150, Flt3, CD16/32 and CD34 Abs. For lymphoid differentiation assays, MPP1, MPP3 or MPP4 (0.5 x 10^3^ cells/well in 96-well plates) were isolated from BM or sorted from MPP1 cultures into wells pre-seeded with 10^5^ OP9 stromal cells. As previously described (10), cells were cultured in OptiMEM (Invitrogen) supplemented with 5% FBS, 1% penicillin/streptomycin, and the following murine cytokines: SCF (10 ng/mL), Flt3-L (10 ng/mL) and IL-7 (5 ng/mL) in the presence or absence of 10 µM AMD3100 or 250 nM Rapamycin. After sequential withdrawal of Flt3-L (day 3) and SCF (day 5), cells were maintained with IL-7 until 7 days of culture. Cells were then harvested and analyzed after by flow cytometry after staining with CD11b and B220 Abs. For Seahorse metabolic flux experiments, OCR and ECAR were measured using an 8-well Seahorse XFp Analyzer according to manufacturer’s instructions (Agilent Technologies). In brief, MPP4 (7 x 10^4^ cells per well) were sorted directly into 8-well plates pre-coated with poly-lysine (Sigma-Aldrich) and containing XF DMEM medium. Plates were then centrifuged and immediately analyzed following manufacturer’s instructions. For measurement of glucose uptake and ROS levels, 2 x 10^3^ purified MPP4 were seeded into a 96-well tissue culture plate in DMEM medium supplemented with 10% FBS, 1% penicillin/streptomycin. After 4 h of culture, cells were washed with DMEM and incubated for 20 min at 37°C with MitoSox (5 µM, Thermo Fisher Scientific), CellRox DeepRed (1.25 µM, Thermo Fisher Scientific) and dichlorofluorescin diacetate (DCFDA, 10 µM, Thermo Fisher Scientific) in order to measure ROS levels by flow cytometry. For glucose uptake experiments, the medium was supplemented with 100 µM of 2- NBD-Glucose (Abcam) at the start of the assay and the incorporation of this fluorescent analog has been measured by flow cytometry after 4 and 20 h of culture.

### *In situ* RNA hybridization and immunofluorescence microscopy

For analyses of BM localization of MPP4 by *in situ* RNA hybridization and immunofluorescence microscopy, freshly dissected mouse femurs were fixed overnight in 4% ParaFormAldehyde (PFA) followed by 48 h decalcification in EDTA (0.5 M, pH=7.4) under agitation at 4°C. Bones were incubated overnight in PBS 1X containing 20% sucrose (Sigma) and 2% polyvinylpyrrolidone (PVP) (Sigma) at 4°C. The embedding was performed using Sub Xero freezing media (Mercedes Medical) before storing the samples at -80°C. Bones were sectioned (14 µm-thick) using a cryostat (Thermofisher). RNA-Scope was performed using the RNAscope® Multiplex Fluorescent Detection Kit v2 kit (323110, ACD) and RNAscope® H_2_O_2_ and protease Reagents kit (322381, ACD) according to the manufacturer’s instructions. Briefly, tissue sections were rehydrated in PBS 1X for 5 min at RT, incubated 30 min at 60°C in a HybEZ II oven and fixed for 15 min at 4°C in 4% PFA. Tissue sections were treated with H_2_0_2_ for 10 min at RT followed by incubation in target retrieval reagents solution for 11 min at 90°C and protease III solution for 30 min at 40°C. Then, sections were incubated with targeted probes: Flt3 (487861 RNAscope® Probe - Mm-Flt3 - musculus FMS-like tyrosine kinase 3 (Flt3) mRNA), c-Kit (314151-C2 RNAscope® Probe Mm-Kit-C2), positive probe (LOT: 2018452 RNAscope® 3-plex positive control probe-Mm) and negative probe (LOT: 21285A RNAscope® 3-plex negative control Probe). The hybridization procedure was performed for 2 h at 40°C. Sequential amplification steps were performed according to manufacturer’s instructions using Amp1, Amp2 and Amp3 solutions at 40°C. Finally, tissue sections were incubated with Opal570 (OP-001003 Bio-thechne R&D) and Opal650 (OP-001005 Bio- thechne R&D) for 30 min at 40°C. RNA *in situ* hybridization was followed by an immunofluorescence overnight staining using the primary anti-mSca-1/Ly6 Ab (AF1226 Bio- thechne R&D). Laminin positive structures were identified by an immunofluorescence overnight staining using the primary anti-Laminin Ab (L9393 Sigma). Then, sections were incubated with the appropriate secondary Abs Donkey anti-goat AF488 (A-11055 Thermofisher) and Donkey anti-Rabbit IgG (H+L) Highly Cross-Adsorbed Secondary Antibody, AF680 (A-10043 Thermofisher) for 1 h 15 min at RT with DAPI for nuclear staining. Autofluorescence was removed with the True Black Kit (92401 TrueBlack® Lipofuscin Autofluorescence Quencher) according to the manufacturer’s instructions. Mounting was performed using the mounting media ProLong Gold antifade reagent (P36934 Invitrogen). Images were acquired using a LSM780 confocal microscope and processed using the open- source digital image analysis software QuPath v0.2.3 (78). Integration of ImageJ (79) package allowed the total cell detection. Sca-1^+^ cell detection and classification were assessed by using the Qupath detection and classification tools. Subcellular detection was performed within the Sca-1^+^ cells to classify MPP4 as Sca-1^+^, Flt3^+^, c-Kit^+^ cells. Laminin detection was performed by pixel classification tool. Distance was calculated from each cell classified as a MPP4 cell to the nearest Laminin^+^, Sca-1^+^ cells by spatial analysis in Qupath using the detectionToAnnotationDistances tool. Ripley’s L statistics was used to detect clustering patterns in point patterns obtained from coordinates of cells detected with the Qupath software. Ripley’s L function is defined as the cumulative distribution function of the nearest-neighbor distances and was computed using the spatstat package in the R statistical software (80). It measures the expected number of points within a certain distance of another point, compared to a random distribution. The L function is defined as: L(d) = √(A/π) - d * Σ(w_ij / n) where L(d) is the value of Ripley’s L function at distance d, A is the total area of the study region, w_ij is the weight associated with the pair of points i and j, n is the total number of points, and the summation is performed over all pairs of points within distance d. By comparing the observed L(d) values with the expected values under a random distribution, it is possible to determine whether the points are clustered, dispersed, or randomly distributed. Observed L(d) values that are greater than the expected values for all distances d indicate clustering of points. On the other hand, if the observed values are consistently lower than the expected values, it suggests a dispersion pattern. If the observed and expected values are similar, it indicates a random distribution.

For mitochondria network analysis by immunofluorescence, 5 x 10^3^ MPP4 were sorted directly into 24-well plates containing coverslips pre-coated with poly-l-lysine. After 2 h incubation at 37°C in order to allow cell adhesion to the coverslips, the cells were fixed using PBS 4% PFA. Fixed cells were permeabilized with PBS 0.3% Triton X for 10 min, extensively washed and blocked with PBS 1% BSA for 1 h at RT. The MPP4 were then incubated with unlabeled primary TOMM20 (1 µg/mL, Rabbit IgG, Thermo Fisher Scientific) Ab overnight at 4°C, followed by wash and incubation with secondary Ab (1 µg/mL, Goat anti-Rabbit IgG, Thermo Fisher Scientific) and the nuclear dye Hoechst 33342 during 1 h at RT (Table S8). After mounting the slides using ProLong™ Gold Antifade mounting medium (Thermo Fisher Scientific), the images were obtained with a Plan-Apochromatic objective (63x oil-immersion objective with a numerical aperture of 1.4) using the LSM800 confocal microscope (Carl Zeiss). Sections were acquired as serial z stacks (0.56 µm apart) and were subjected to maximum intensity projection in the Zen 2.3 System software. Brightness and contrast settings were set during capture and not altered for analysis. Mitochondrial labeling were individualized for each confocal optical slice and the number of fragments were calculated using the ‘Analyze Particles’ plug-in on thresholded images in Fiji software using a threshold of 4483 and a filter of 0.6µm^2^ for the minimum size of the fragments. A macro enabling the automatic quantification of a batch of images is available upon request. At least 50 cells/condition were analyzed.

### Statistical analyses

Data are expressed as mean ± SEM. All statistical analyses were conducted using Prism software (GraphPad Software). A Kruskal-Wallis test was used to determine the significance of the difference between means of WT, +/1013 and 1013/1013 groups (^#^*P* < 0.05; ^##^*P* < 0.005; and ^###^*P* < 0.0005). The unpaired two-tailed Student *t* test was used to compare means among two groups. The non-parametric Wilcoxon–Mann–Whitney test was used in mitochondria network analyses to determine the statistical differences between the median of two groups.

## LIST OF SUPPLEMENTARY MATERIALS

**Table S1: Complete GO analysis of genes associated with down ATAC-seq peaks in mutant MPP1 related to Figure 2**.

**Table S2: List of HOMER known motifs down in mutant MPP1 related to Figure 2**.

**Table S3: Complete GO analysis of genes associated with down ATAC-seq peaks in mutant MPP4 related to Figure 2**.

**Table S4: List of HOMER known motifs down in mutant MPP4 related to Figure 2**.

**Table S5: Complete GO analysis of genes up- and down-regulated in mutant MPP4related to Figure 4**.

**Table S6: Biomark principal component analysis contributing genes related to Figure 5**.

**Table S7: List of primers used for the BioMark assay related to the main Figure 5 and** SF1.

<u>Table S8:</u> List of antibodies used for immunofluorescence related to the main Figure 6.

**Figure S1 related to Figure 1**

**Figure S2 related to Figure 2**

**Figure S3 related to Figure 3**

**Figure S4 related to Figure 4**

**Figure S5 related to Figure 5**

**Figure S6 related to Figure 6**

**Figure S7 related to Figure 7**

## ACKNOWLEDGMENTS

We thank Drs. C. Doliger and S. Duchez (Flow Cytometry Core Facilities, Institut de Recherche Saint-Louis, Paris) for their technical assistance. The authors gratefully acknowledge the contribution of *Cxcl12^DsRed^* knock-in reporter mice generated by S.J. Morrison (Ding and Morrison, 2013) and provided by Dr. L.G. Ng (Singapore Immunology Network, A*STAR, Singapore). We thank Dr. C. Jones and C. O’Brien (Princess Margaret Cancer Centre, Toronto, Canada) for editing the writing of our manuscript. The study was supported by the Laboratory of Excellence in Research on Medication and Innovative Therapeutics (LabEx LERMIT) (ME and KB), an ANR JCJC grant (ANR-19-CE15-0019-01) to ME and an ANR PRC grant (ANR- 17-CE14-0019), an INCa grant (PRT-K 2017), the Association Saint Louis pour la Recherche sur les Leucémies and the FRM (Programme Equipe FRM 2022, EQU202203014627) to KB.V.R. was supported by the FRM, La Ligue Contre le Cancer and la Société Française d’Hématologie. M.K. was fellowship recipient from the French Ministry. Z.A-N. was fellowship recipient from the French Ministry and then supported by the FRM. J.L. was recipient from the People Programme (Marie Curie Actions) of the European Union’s Seventh Framework Programme (FP7/2007-2013) under REA grant agreement n. PCOFUND-GA- 2013-609102, through the PRESTIGE programme coordinated by Campus France, and from a ANR grant (ANR-17-CE14-0019). D.H.M and P.M.M. were supported by the Division of Intramural Research of the National Institute of Allergy and Infectious Diseases, National Institutes of Health.

## AUTHOR CONTRIBUTIONS

V.R. designed and performed most of the experiments and contributed to manuscript writing; Ma.K., L.R., Z.A.N., V.G., A.B., J.L., Mé.K., C.M., B.S., S.L., C.F. and A.A. performed some of the experiments, analyzed data and reviewed the manuscript; M.D. and G.B. analyzed RNA-seq raw data; N.M., N.S., N.D., F.B. and M.A-L. contributed to data analyses and reviewed the manuscript; M.E. and S.J.C.M. helped with the study design, performed some of the experiments, contributed to data analyses and reviewed the manuscript; V.P. performed intravenous injections during *in vivo* experiments and reviewed the manuscript; D.S. performed artificial intelligence-based image analyses and reviewed the manuscript; C.L. performed and analyzed epigenetic studies and reviewed the manuscript; D.H.M. and P.M.M provided WS samples and clinical data and reviewed the manuscript; K.B. conceived, designed and supervised the study, contributed to data analyses, found funding for the study, and wrote the manuscript.

