## Supplemental Information for "CXCR4 signaling strength regulates hematopoietic multipotent progenitor fate through extrinsic and intrinsic mechanisms"

### LIST OF SUPPLEMENTARY MATERIALS

#### SUPPLEMENTAL TABLES

**Table S1:** Complete GO analysis of genes associated with down ATAC-seq peaks in mutant MPP1 related to Figure 2.

**Table S2:** List of HOMER known motifs down in mutant MPP1 related to Figure 2.

**Table S3:** Complete GO analysis of genes associated with down ATAC-seq peaks in mutant MPP4 related to Figure 2.

**Table S4:** List of HOMER known motifs down in mutant MPP4 related to Figure 2.

**Table S5:** Complete GO analysis of genes up- and down-regulated in mutant MPP4 related to Figure 4.

**Table S6:** Biomark principal component analysis contributing genes related to Figure 5.

**Table S7:** List of primers used for the BioMark assay related to the main Figure 5 and SF1.

**Table S8:** List of antibodies used for immunofluorescence related to the main Figure 6.

#### SUPPLEMENTAL FIGURES

**Figure S1 related to Figure 1**

**Figure S2 related to Figure 2**

**Figure S3 related to Figure 3**

**Figure S4 related to Figure 4**

**Figure S5 related to Figure 5**

**Figure S6 related to Figure 6**

**Figure S7 related to Figure 7**

**SUPPLEMENTAL TABLES**

**Table S7: List of primers used for the BioMark assay related to the main Figure 5 and SF1.**

| Genes | Primers |
|---|---|
| **MPP markers** | |
| *Cd48* | Mm00455932_ml |
| *Slamf1* | Mm00443316_ml |
| *Flt3* | Mm00439016_ml |
| *Irf8* | Mm00492567_ml |
| **Migration** | |
| *Cxcl12* | Mm00445553_ml |
| *Cxcr4* | Mm01292123_ml |
| *Ackr3* | Mm01290382_ml |
| **Lymphomyeloid differentiation** | |
| *Ikzf1* | Mm01187880_ml |
| *Il7r* | Mm00434295_ml |
| *Dntt* | Mm00493500_ml |
| *Bach2* | Mm00464379_ml |
| *Mpo* | Mm00447886_ml |
| *Cebpa* | Mm00514283_s1 |
| *Mpl* | Mm00440310_ml |
| *Gfi1b* | Mm00492318_ml |
| **Irrelevant genes (negative controls)** | |
| *Pax5* | Mm00435501_ml |
| *Cd3e* | Mm00599684_g1 |
| **Housekeeping gene** | |
| *Actb* | Mm01205647_g1 |

**Table S8: List of antibodies used for immunofluorescence related to the main Figure 6.**

| Antibody | Origin | Conjugate | Supplier |
|---|---|---|---|
| Tomm20 | Rabbit | Uncoupled | Thermo Fischer Scientific |
| IgG (Secondary Antibody) | Goat | Alexa Fluor 555 | Thermo Fischer Scientific |

### SUPPLEMENTAL FIGURES

**Figure S1 related to Figure 1:** **(A)** Expression levels of *Cxcr4*, *Ackr3* and *Cxcl12* were determined by quantitative real-time PCR in MPP2, MPP3 and MPP4 sorted from the marrow fraction of WT and mutant mice. Relative expression levels (RQ) have been normalized with *Actb* expression levels in each sample (6-12 pools of 0.1 x $10^3$ cells *per* condition). Data are from two independent experiments and each individual sample was run in duplicate. **(B)** Expression levels of Cxcr4 were determined by flow cytometry on gated MPP2, MPP3 and MPP4 from the marrow fraction of WT and mutant mice. Cxcr4-positive fractions (left) or MFI values (right) obtained within MPP2, MPP3 and MPP4 relative to background fluorescence based on the corresponding isotype control staining are shown. Data are from at least four independent experiments with 8-11 mice per group. **(C)** Left: Representative histograms for surface detection of Ackr3 on gated MPP4 and $Lin^+$ cells ($CD3^+B220^+Gr1^+Ter119^+CD41^+$) from marrow fraction of WT and mutant mice. Background fluorescence is shown (isotype, gray histogram). Right: MFI values for Ackr3 were determined by flow cytometry on gated $Lin^+$ cells, MPP2, MPP3 and MPP4 from the marrow fraction of WT and mutant mice. Data are from two independent experiments with 3 mice per group. **(D)** Left: Representative flow cytometric analyses for GFP expression in MPP4 or $Lin^+$ cells from marrow fraction of $\textit{Ackr3}^{GFP}$ reporter mice. Right: Proportions of $GFP^+$ cells on gated $Lin^+$ cells, MPP2, MPP3 and MPP4 from the marrow fraction of $\textit{Ackr3}^{GFP}$ reporter mice. **(E)** Left: Representative flow cytometric analysis for Dsred expression in MPP4 from marrow fraction of $\textit{Cxcl12}^{Dsred}$ reporter mice. Right: Proportions of $Dsred^+$ cells on gated MPP1, MPP2, MPP3 and MPP4 from the marrow fraction of $\textit{Cxcr4}^{+/+}\textit{Cxcl12}^{Dsred}$, $\textit{Cxcr4}^{+/1013}\textit{Cxcl12}^{Dsred}$ or $\textit{Cxcr4}^{1013/1013}\textit{Cxcl12}^{Dsred}$ mice. **(F)** Cell surface expression of Cxcr4 on WT and mutant MPP2/3 or MPP4 upon exposure to 10 nM Cxcl12 at 37°C or 4°C for 45 min. Cxcr4 expression on MPP2/3 or MPP4 incubated in medium alone was set at 100%. Data are summarized from three independent experiments with 5-10 mice per group. **(G)** Absolute numbers of T cells ($CD3^+$), B cells ($B220^+$) and neutrophils ($Gr1^+$) in the blood of WT or mutant mice. Data are from two independent experiments with 6 mice per group. **(H)** Upper: Representative dot-plots comparing the frequencies of HSCs/MPPs ($CD45RA^-CD38^-CD34^+$), MLPs ($CD45RA^+CD38^-CD7^-CD34^+CD10^+$), GMPs ($CD45RA^+CD38^+CD7^-CD34^+CD10^-CD135^+$), CMPs ($CD45RA^-CD38^+CD7^-CD34^+CD10^-CD135^+$) and MEPs ($CD45RA^-CD38^+CD7^-CD34^+CD10^-CD135^-$). Lower: Absolute numbers of HSCs/MPPs, MLPs, GMPs, CMPs and MEPs in BM aspirates of HD and WS patients. Horizontal lines correspond to median values. All displayed results are represented as means ± SEM. Kruskal–Wallis H test–associated p-values (#) are indicated. **, P < 0.005; and ***, P < 0.0005 compared with WT cells; §, P < 0.05 compared with +/1013 cells (as determined using the two-tailed Student's t test).

**Figure S2 related to Figure 2:** **(A)** RNAseq data were generated from 3 x $10^3$ MPP1-4 sorted from the marrow fraction of WT and mutant mice. **(B)** Left: Representative flow cytometric analyses comparing the frequencies of apoptotic (Annexin-$V^+DAPI^-$) MPP3 in WT and 1013/1013 mice. Right: The proportions of apoptotic (Annexin-$V^+DAPI^-$) MPP2, MPP3 and MPP4 were determined by flow cytometry in the marrow fraction of WT and mutant mice. Data are from four independent experiments with 8-12 mice per group. **(C)** Ki-67 and DAPI co-staining was used to analyze by flow cytometry the cell cycle status of MPP1-4 subsets in the marrow fraction of WT and mutant mice. Representative flow cytometric analyses comparing the frequencies of MPP3 and MPP4 in the different phases of the cell cycle in WT and mutant mice are shown. Data are from five independent experiments with 12-15 mice per group. **(D)** Left: Representative flow cytometric analyses comparing the frequencies of $BrdU^+$ MPP4 in WT and mutant mice after a 12-days pulse period. Right: Percentage of $BrdU^+$ cells among WT and mutant MPP2, MPP3 and MPP4 after a 12-days pulse period (left) or 3 weeks of chase (right). Data are from 2 independent experiments with 4-6 mice per group. All displayed results are represented as means ± SEM. Kruskal–Wallis H test–associated p-values (#) are indicated. *, $P < 0.05$; and ***, $P < 0.0005$ compared with WT cells; §, $P < 0.05$ compared with +/1013 cells (as determined using the two-tailed Student's t test).

**Figure S3 related to Figure 3:** **(A)** Representative BM sections from WT mice after targeting Flt3 (i) and c-Kit (ii) by RNA-scope and Sca-1 (iii) and Laminin (iv) by immunofluorescence in association with DAPI. The color code for markers used is indicated and image obtained after merging signals corresponding to the different markers is shown (v). MPP4 defined as c-$Kit^+Sca\text{-}1^+Flt3^+$ are highlighted in yellow and Laminin areas in light blue (vi). Scale bars denote 20 µm. **(B)** Analysis of the spatial distribution of MPP4 cells using Ripley's L statistics defined as the cumulative distribution function of the nearest-neighbor distances (d). It measures the level of clustering or dispersion patterns from point coordinates. The green, red and blue lines indicate respectively the L-functions for the WT, +/1013 and 1013/1013 mice. The grey area shows the simulated values obtained with a complete spatial randomness null model. Among the six BM analyses (two per groups), two showed a partial distribution pattern significantly higher (WT and 1013/1013) from the random pattern demonstrating a partial clustered pattern. **(C)** Representative BM section from WT mice showing arteriolar cells defined as $Lam^+Sca\text{-}1^+$ and highlighted in purple. **(D)** Proportions of arteriolar cells ($Lam^+Sca\text{-}1^+$) detected by immunofluorescence in the BM sections of WT and mutant mice. Data are from 3 independent determinations with 3 mice per group. All displayed results are represented as means ± SEM.

**Figure S4 related to Figure 4:** **(A)** PCA of RNAseq gene expression data obtained from the indicated MPP subpopulations in WT and mutant mice. **(B)** GSEA for MPP2-, MPP3- and MPP4-gene specific signatures between the indicated WT subpopulations. **(C)** Left: Volcano plot of differentially expressed genes in +/1013 (left) vs. WT (right) MPP3. Right: Volcano plot of differentially expressed genes in 1013/1013 (left) vs. WT (right) MPP3. Genes highly differentially expressed (log2FC(mutant vs WT MPP3) > 2; -log10(adjusted p-value) > 2) are shown in red.

**Figure S5 related to Figure 5:** **(A)** *In vivo* differentiation assays workflow. Equal numbers ($10^4$ cells) of sorted CD45.2$^+$ WT or mutant MPP4 were transplanted into sub-lethally irradiated WT CD45.1$^+$ recipient mice and multi-lineage output was determined 2 weeks after transplantation. **(B)** Proportions of donor CD45.2$^+$ (WT, +/1013, or 1013/1013) myeloid cells (Gr1$^+$B220$^-$) recovered from the blood of CD45.1$^+$ recipients two weeks after the transplantation. All displayed results are represented as means ± SEM. Kruskal–Wallis H test–associated p-values (#) are indicated. **, P < 0.005; and ***, P < 0.0005 compared with WT cells (as determined using the two-tailed Student's t test).

**Figure S6 related to Figure 6:** **(A)** Left: RNAseq-based GSEA for the Oxphos signature in +/1013 vs. WT MPP4 and in 1013/1013 vs. WT MPP4. Right: Heatmaps showing normalized dye intensity for the expression of genes involved in the Oxphos signature in WT vs mutant MPP4. **(B)** RNAseq-based GSEA for glycolysis signatures in WT and mutant MPP4. **(C)** WT and mutant MPP4 were stimulated or not with 20 nM Cxcl12 at 37°C for 2 min. Left: Representative histograms for intracellular detection of phospho-p38 on sorted MPP4 from the marrow fraction of WT and mutant mice. Fluorescence in absence of Cxcl12 stimulation is shown (gray histogram). Right: MFI values for phospho-p38 were determined by flow cytometry on sorted MPP4 from the marrow fraction of WT and mutant mice. Data are from two independent experiments with 2-3 mice per group. **(D)** Left: Representative histograms for intracellular detection of ROS by flow cytometry in WT and mutant MPP4. Right: Bar graphs denote the fold change in MitoSox, CellRoxDeepRed and DCFDA MFI values compared with the levels obtained in WT MPP4. Data are from three independent experiments with 5-7 mice per group. **(E)** Glucose uptake in WT and mutant MPP4 at the start of the assay or after 4 or 20 hours of culture. MFI values for 2-NBDG have been determined by flow cytometry. Data are from three independent experiments with 4-6 mice per group. **(F)** Left: Representative histograms for TFAM staining in gated MPP4 from marrow fraction of WT and mutant mice. Right: MFI values of TFAM in WT and mutant MPP4. Data are from two independent experiments with 3 mice per group. **(G)** WT and mutant total BM cells pre-incubated or not with 10 μM of AMD3100 were stimulated or not with 20 nM Cxcl12 at 37°C for 5 min. MPP1 were then stained with specific Abs. Representative histograms for intracellular detection of phospho-mTOR and phospho-S6 on gated MPP1 from the marrow fraction of WT and mutant mice are shown. Fluorescence in absence of Cxcl12 stimulation (gray histogram) or presence of Cxcl12 and AMD3100 (black histogram) are displayed. MFI values for phospho-mTOR and phospho-S6 were determined by flow cytometry. Data are from three independent experiments with 4-6 mice per group. **(H)** MFI values for TMRE and MTG in WT and mutant MPP1. Data are from three independent experiments with 9-10 mice per group. **(I)** Left: Representative histograms for TFAM staining in gated MPP1 from marrow fraction of WT and mutant mice. Right: MFI values of TFAM in WT and mutant MPP1. Data are from two independent experiments with 3 mice per group. *, $P < 0.05$; **, $P < 0.005$; and ***, $P < 0.0005$ compared with WT cells; $^{\Delta}$, $P < 0.05$; $^{\Delta\Delta}$, $P < 0.005$; and $^{\Delta\Delta\Delta}$, $P < 0.0005$ compared with non-stimulated cells.

**Figure S7 related to Figure 7:** **(A and B)** Equal numbers of the fraction B (TMRE$^{low}$) sorted from WT or mutant MPP4 were cultured for 4 or 7 days in a cytokine-supplemented feeder-free media in the presence or absence of 10 μM AMD3100 or 250 nM Rapamycin. Viability

(A) and proportions of MPP4 (B) were assessed at day 4 by flow cytometry. Data are from two independent experiments with 3-4 mice per group. [£], $P < 0.05$; and [££], $P < 0.005$ compared with PBS-treated cells (as determined using the two-tailed Student's t test).

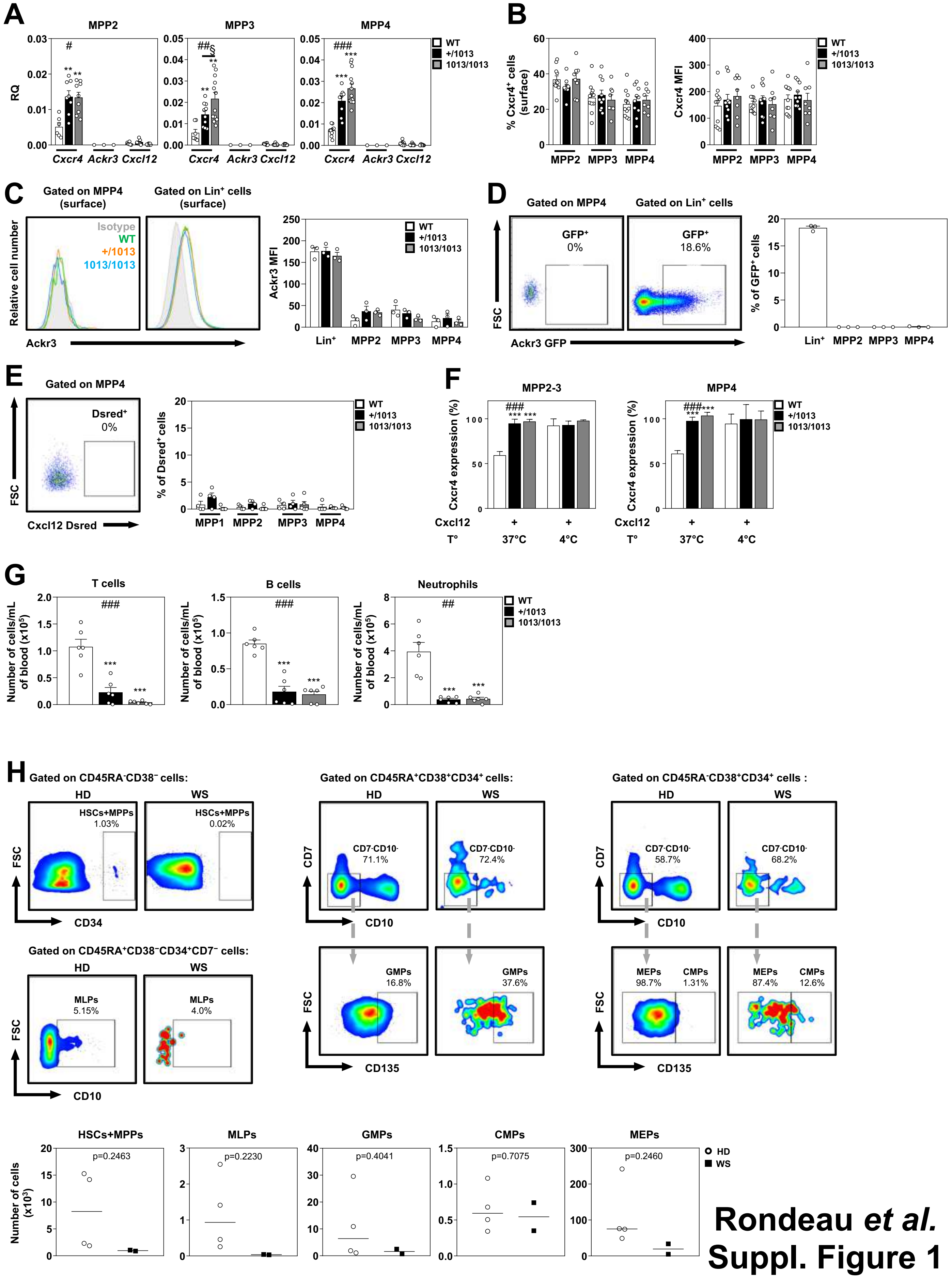


**Rondeau *et al.* Suppl. Figure 1**

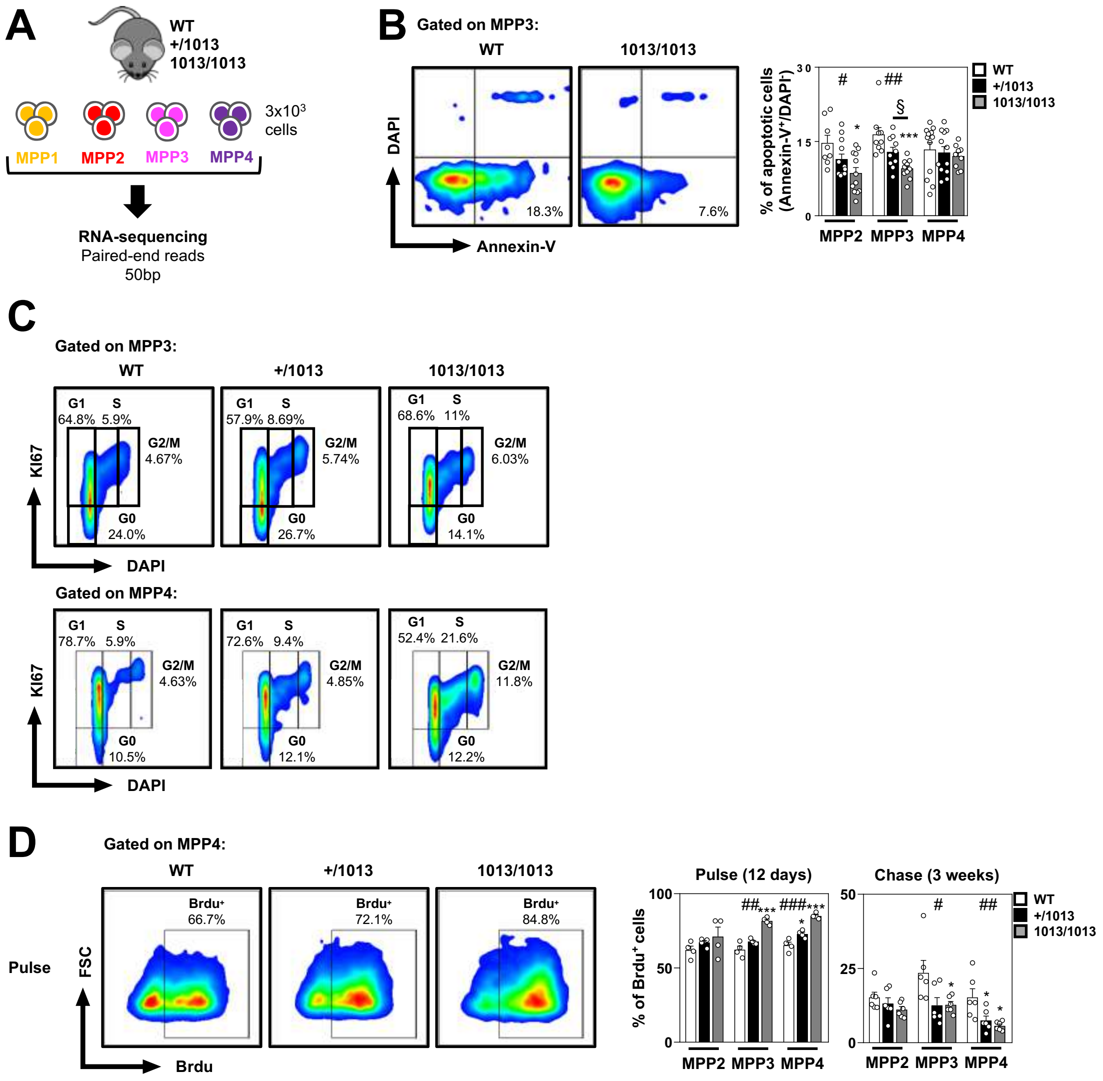


Rondeau *et al.*
Suppl. Figure 2

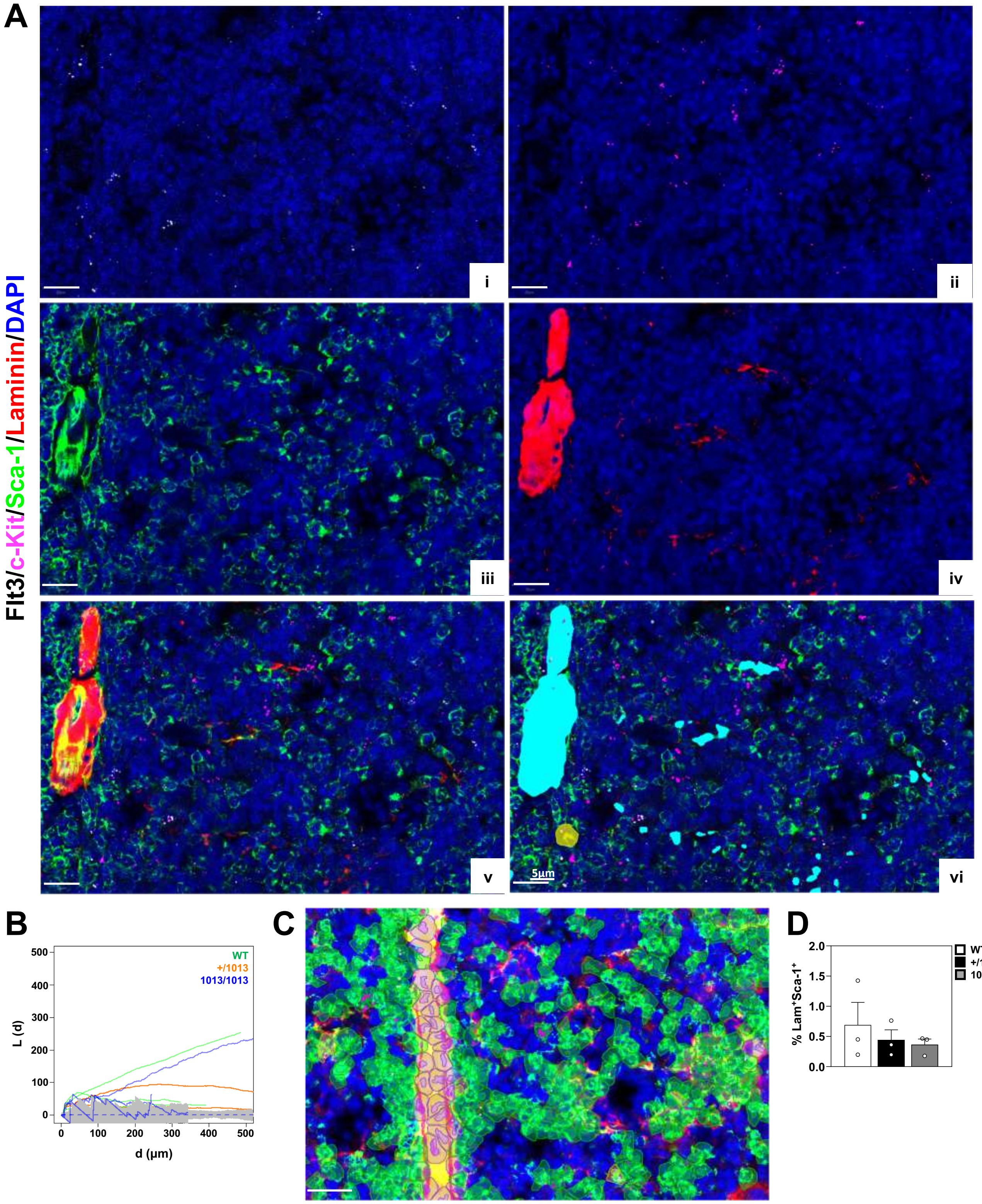


**Rondeau *et al.***
**Suppl. Figure 3**

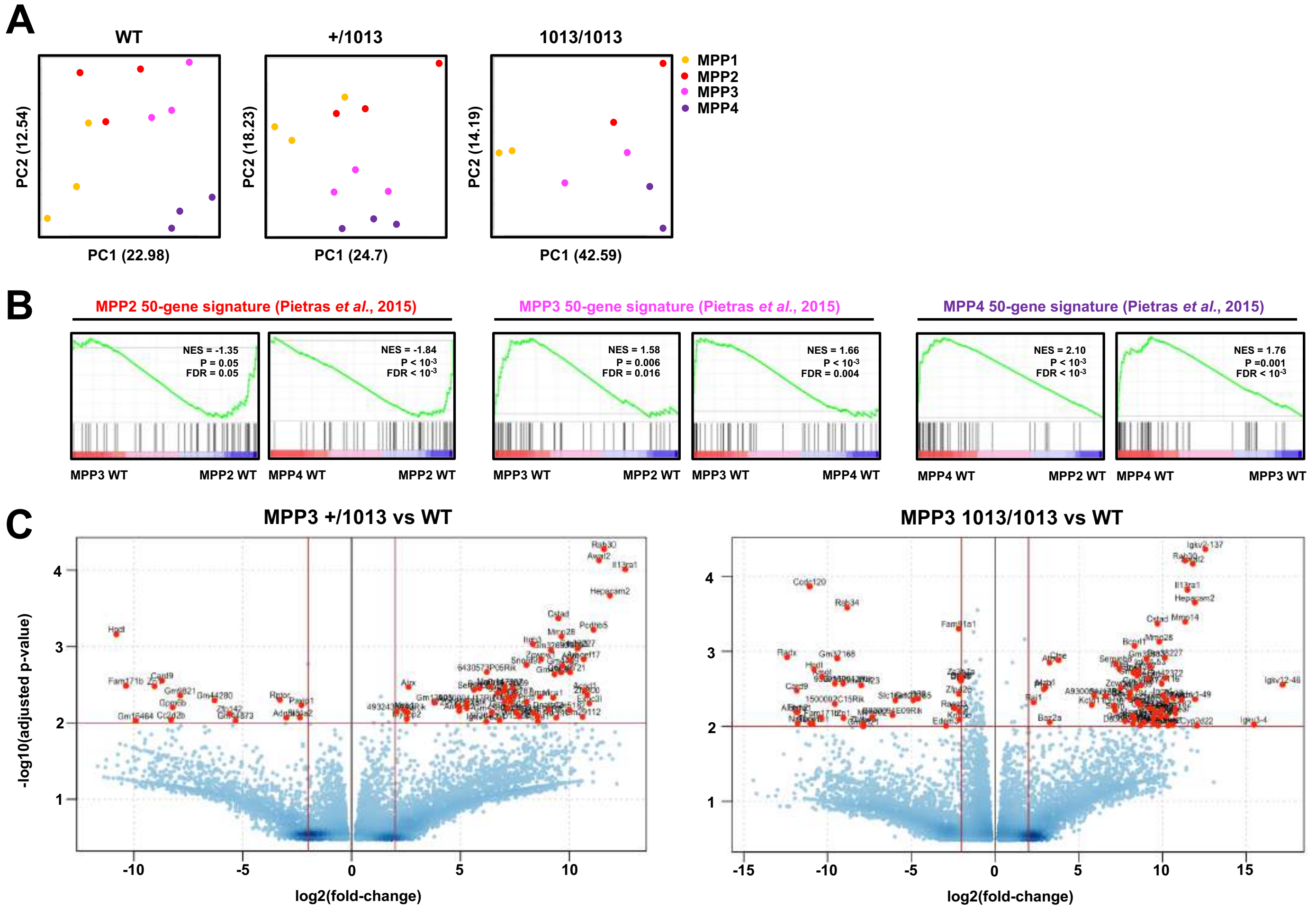


**Rondeau *et al.***
**Suppl. Figure 4**

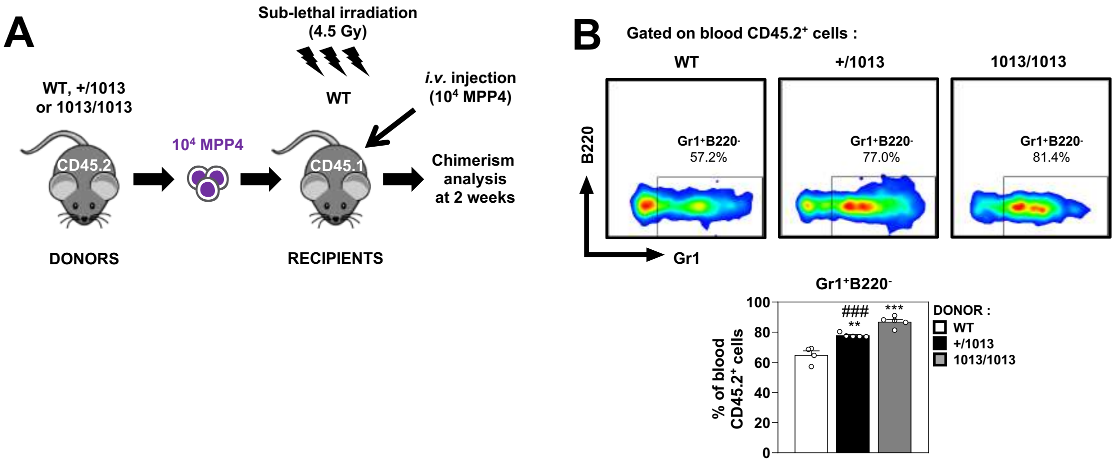


Rondeau *et al.*
Suppl. Figure 5

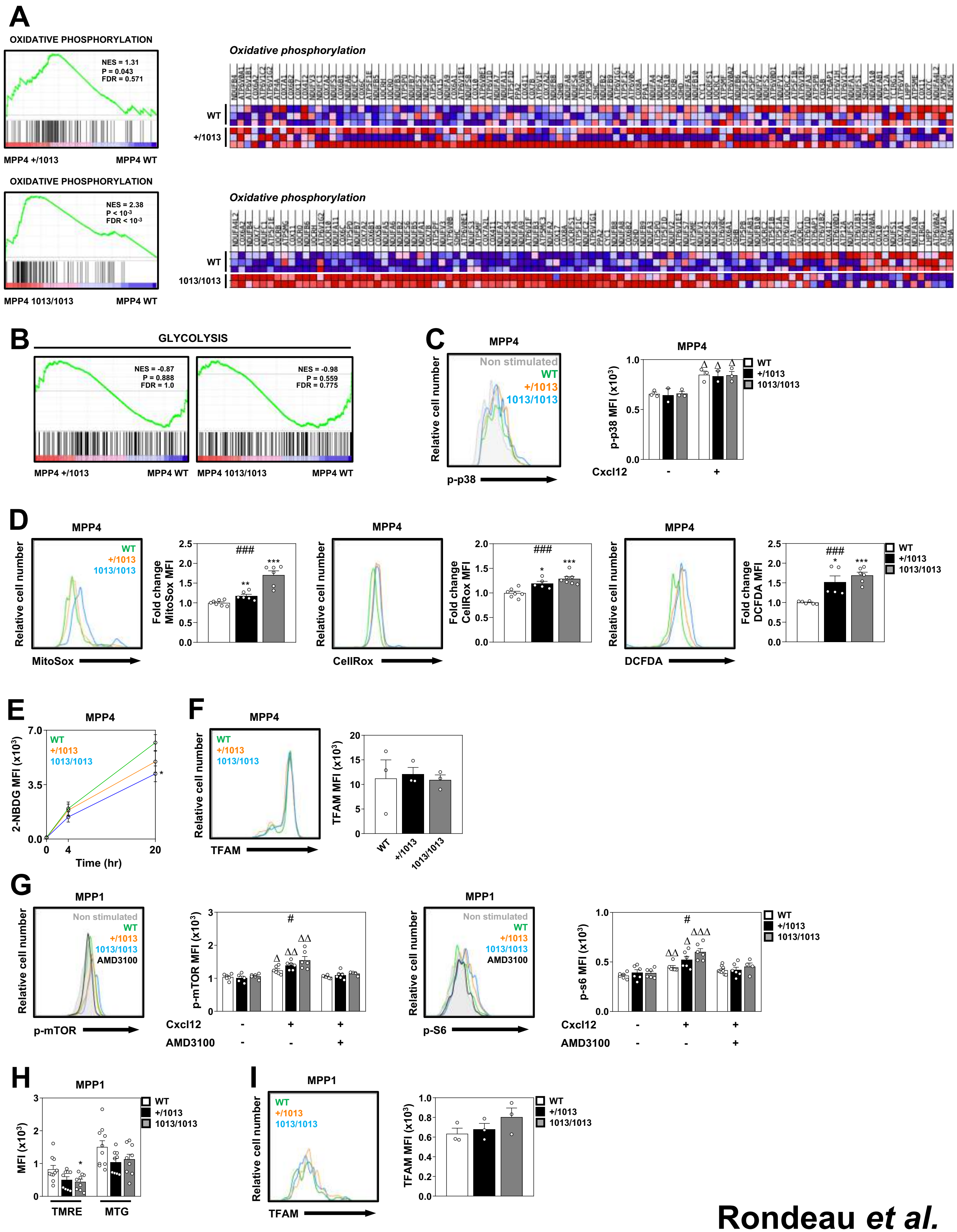


**Rondeau *et al.* Suppl. Figure 6**

A

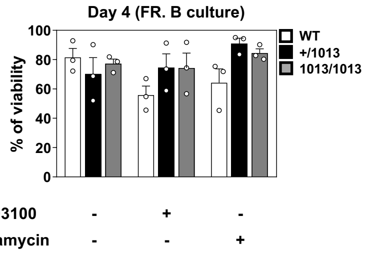


B

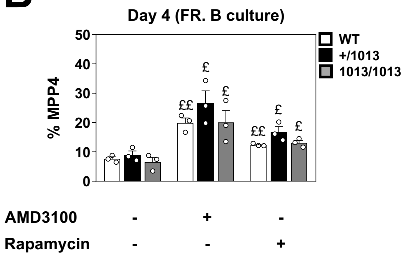


Rondeau *et al.*
Suppl. Figure 7
